# PCM1 conveys centrosome asymmetry to polarized endosome dynamics in regulating daughter cell fate

**DOI:** 10.1101/2024.06.17.599416

**Authors:** Xiang Zhao, Yiqi Wang, Vincent Mouilleau, Ahmet Can Solak, Jason Garcia, Xingye Chen, Christopher J. Wilkinson, Loic Royer, Zhiqiang Dong, Su Guo

**Affiliations:** Chan Zuckerberg Biohub, San Francisco, California, 94158, USA; Department of Bioengineering and Therapeutic Sciences, Programs in Biological Sciences and Quantitative Biosciences, Institute of Human Genetics, Kavli Institute for Fundamental Neuroscience, Bakar Aging Research Institute, University of California, San Francisco, California, 94158, USA; Lawrence Livermore National Laboratory, Livermore, CA 94550, USA; Department of Automation, Tsinghua University, Beijing, China; Centre for Biomedical Sciences, Department of Biological Sciences, Royal Holloway University of London, Egham, Surrey, TW20 0EX, UK; College of Biomedicine and Health, College of Life Science and Technology, Huazhong Agricultural University, Wuhan 430070, China

**Keywords:** neurogenesis, cancer, neuropsychiatric disorders, Delta D, Notch, neural rosettes, brain organoids, hiPSCs

## Abstract

Vertebrate radial glia progenitors (RGPs), the principal neural stem cells, balance self-renewal and differentiation through asymmetric cell division (ACD), during which unequal inheritance of centrosomes is observed. Mechanistically, how centrosome asymmetry leads to distinct daughter cell fate remains largely unknown. Here we find that the centrosome protein Pericentriolar Material 1 (Pcm1), asymmetrically distributed at the centrosomes, regulates polarized endosome dynamics and RGP fate. *In vivo* time-lapse imaging and nanoscale-resolution expansion microscopy of zebrafish embryonic RGPs detect Pcm1 on Notch ligand-containing endosomes, in a complex with the polarity regulator Par-3 and dynein motor. Loss of *pcm1* disrupts endosome dynamics, with clonal analysis uncovering increased neuronal production at the expense of progenitors. Pcm1 facilitates an exchange of Rab5b (early) for Rab11a (recycling) endosome markers and promotes the formation of Par-3 and dynein macromolecular complexes on recycling endosomes. Finally, in human-induced pluripotent stem cell-derived brain organoids, PCM1 shows asymmetry and co-localization with PARD3 and RAB11A in mitotic neural progenitors. Our data reveal a new mechanism by which centrosome asymmetry is conveyed by Pcm1 to polarize endosome dynamics and Notch signaling in regulating ACD and progenitor fate.

## Main

Vertebrate RGPs are the neural stem cells (NSCs) that undergo orchestrated divisions to produce diverse cell types in the central nervous system ^1–6^. During active neurogenesis in developing vertebrate embryos, most RGPs undergo asymmetric cell division (ACD), resulting in a self-renewing and a differentiating daughter with prominent asymmetry of Notch signaling ^5, 6^. Notch asymmetry between daughter cells is a conserved feature in both invertebrates ^7, 8^ and vertebrates ^5, 6, 9, 10^. Studies in both Drosophila ^7, 8^ and zebrafish ^11, 12^ have shown that Notch asymmetry between daughter cells is laid down in the mother via asymmetric distribution of Notch ligand-containing endosomes. In the zebrafish forebrain RGPs, which mostly divide along the anteroposterior (A-P) embryonic axis, endosomes containing the Notch ligand Delta D (Dld) are enriched toward the posterior side at anaphase. This process is critically dependent on the polarity regulator Par-3 (also known as PARD3 in humans, and Bazooka in Drosophila) and the dynein motor complex, both localized on the endosomes ^12^. How they assemble on the Notch ligand-containing endosomes to drive posterior-directed movement is unclear.

Another organelle that displays asymmetry during RGP division is the centrosome, which undergoes semi-conservative duplication during S-phase, resulting in two centrosomes that differ in age and protein composition ^13^. Following mitosis, the mother centrosome has a higher activity than the daughter in organizing microtubules to grow the primary cilium. Asymmetric inheritance of centrosomes has been associated with distinct daughter cell fate in RGPs ^14^ and in other cell types ^15, 16^, but how centrosome asymmetry confers daughter cell fate differences at the mechanistic level remains unclear.

In animal cells, the centrosome is composed of a pair of centrioles embedded in a cloud of electron-dense pericentriolar material (PCM) ^17–19^, of which Pcm1 is a core component involved in maintaining centrosome integrity, anchoring microtubules ^20, 21^, and ciliogenesis ^22–24^. The human *PCM1* has been implicated in several forms of cancer and neuropsychiatric disorders ^22, 23, 25, 26^, but the *in vivo* dynamics and function of PCM1 in mitotic progenitors are unclear. Here we reveal a previously unknown asymmetry of Pcm1 at the centrosomes and moreover its localization on the endosomes in embryonic zebrafish forebrain RGPs. Using *in vivo* time-lapse imaging, high resolution microscopy, and molecular genetic approaches, we further uncover a critical role of Pcm1 in the asymmetric segregation of recycling Notch ligand-containing endosomes; it does so by facilitating the formation of macromolecular complexes composed of Par-3 and dynein/dynactin on the recycling endosomes. Asymmetry of PCM1 and its co-localization with PARD3 and the recycling endosomal protein RAB11 are also observed in hiPSC-derived neural progenitors.

## Results

### Pcm1 localization to centrosomes and endosomes in zebrafish forebrain mitotic RGPs

Given the importance of centrosomes in organizing RGP division ^14, 27^, we studied the role of Pcm1, a conserved component of the pericentriolar satellites ^20^. Using a custom generated antibody (see Methods), we first examined the subcellular distribution of Pcm1 in mitotic RGPs. During active neurogenesis (24-30 hours post fertilization, hpf), most forebrain RGPs divide along the anteroposterior (A-P) embryonic axis ^12^. At metaphase, Pcm1 was distributed in a cloud-like pattern around the γ-tubulin-positive centrosomes. Asymmetry of the cloud was often detected, and based on its known role in ciliogenesis, Pcm1 is likely enriched around the mother (here, posterior) centrosome that seeds primary cilia growth ^28^. Quantification uncovered that most RGPs had Pcm1 enriched around the posterior centrosome (**Fig. 1a, 1e**). During anaphase and early telophase, the enrichment of Pcm1 around posterior centrosomes increased (**Fig. 1b, 1e**).

Quite unexpectedly, Pcm1 immunoreactivity was frequently observed in between the separating nuclei (**Fig. 1b, 1d**), a region that we referred to as the central zone. The central zone harbors central spindles, where active organelle trafficking and sorting take place. Quantification uncovered variable amounts of Pcm1 in the central zone relative to its total immunoreactivity in the cell (**Fig. 1f**). These variabilities reflect either the dynamics or the heterogeneity of Pcm1 distribution in the central zone.

The specificity of Pcm1 localization patterns was validated using both *pcm1* morphants (MO) and *pcm1* knock-out (KO) embryos derived from homozygous mutant parents generated via CRISPR genome editing (**Fig. 1c**, and **Extended Data Fig. 1a-1c**). Embryonic development was disrupted in both *pcm1* MO and KO embryos **(Extended Data Fig. 1d-f**).

**Fig 1.**
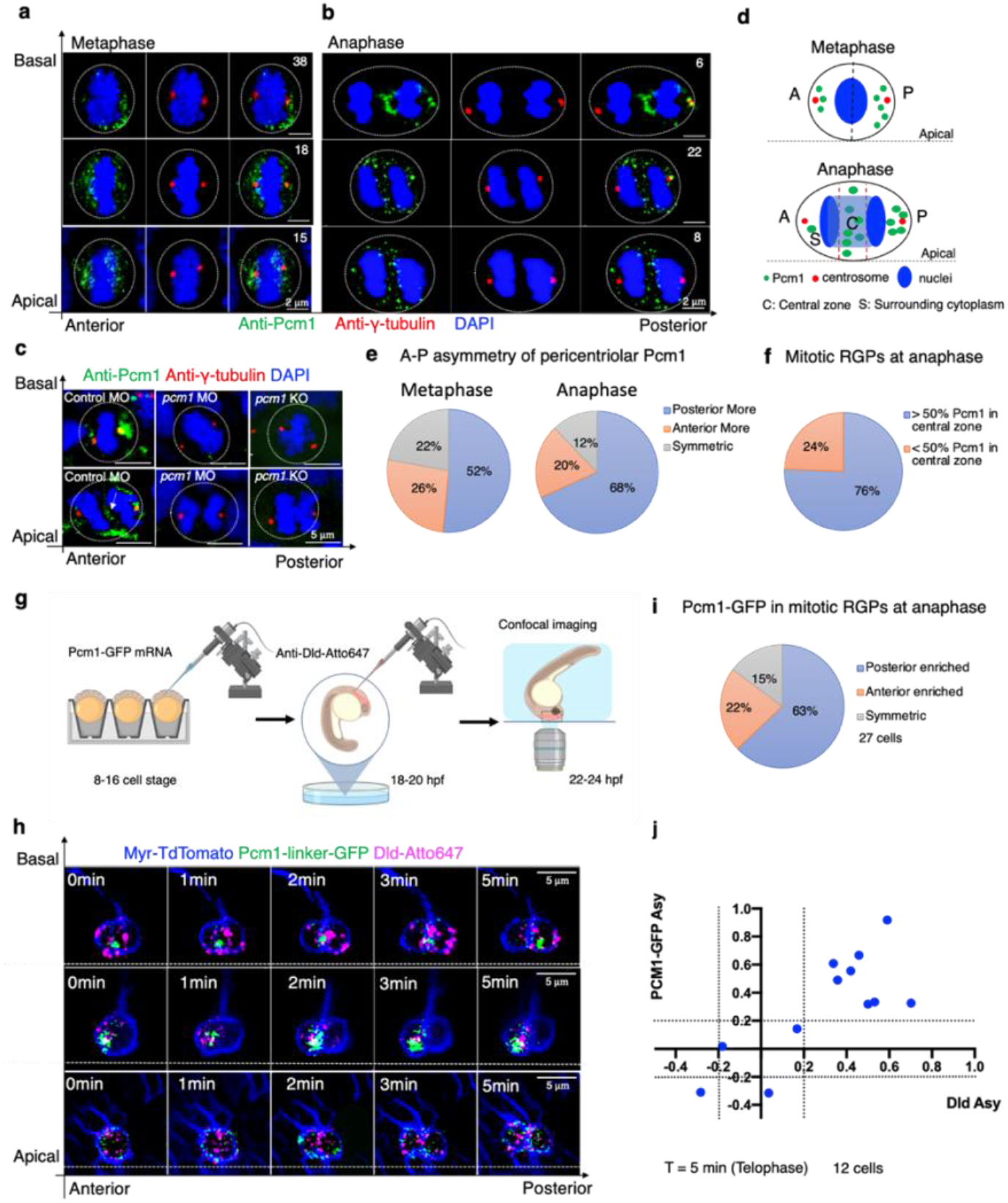
Pcm1 expression is enriched around the posterior centrosome and in the central zone of zebrafish embryonic forebrain mitotic RGPs. **a**. Metaphase RGPs with posterior enrichment (upper row, n=38), anterior enrichment (middle row, n=18) and symmetric (lower row, n=15) peri-centrosome (CEN) localization of Pcm1. **b**. Anaphase RGPs with Pcm1 detected in the central zone (Top: intense labeling, n=6; middle: scattered labeling with posterior CEN enrichment, n=22; bottom, scattered labeling with anterior CEN enrichment, n=8). RGPs were from 5 embryos of 28-32 hpf. Scale bars, 2 μm . **c**. Mitotic RGPs from control MO, *pcm1* MO, and *pcm1* KO embryos. Scale bars, 5 μm . Maximum intensity projection (MIP) of 8-10 confocal z-stacks (0.26 μm z-step) are shown for each image. **d**. Schematic of Pcm1 localization in mitotic RGPs. **e**. Pie charts summarizing the pericentriolar asymmetry of Pcm1 in metaphase (n=68) and anaphase (n=41) RGPs. **f**. Pie chart summarizing the Pcm1 distribution in the central zone of anaphase RGPs (n=41). **g**. Schematic of *in vivo* imaging of Pcm1-GFP. **h**. Three montages show typical Pcm1-GFP dynamics with live labeling of Dld endosomes. In all images, MIP of 5 μm z-stacks (1 μm z-step) are shown. Time is registered using the appearance of cleavage furrow between daughter cells as a landmark. **i**. Pie chart shows statistics of Pcm1-GFP A-P asymmetry at T = 1 min (anaphase). N=27 from 6 embryos. **j**. scatterplot of individual RGP’s asymmetry indices for internalized Dld endosomes (x axis) and Pcm1-GFP (y axis), the dotted lines indicate the asymmetric index threshold: <-0.2: anterior enriched; >0.2: posterior enriched. N=12 from 4 embryos.

Given the prominent localization of Pcm1 in the central zone, we wondered whether it might be associated with Dld endosomes, which are previously shown to converge toward the central zone followed by posterior enrichment ^12^. To observe the *in vivo* dynamics of Pcm1 together with Dld endosomes, we generated GFP-tagged Pcm1 (Pcm1-linker-GFP). Pcm1 is a large protein of over 2000 amino acids with alternatively spliced forms reported in the database (ENSEMBL ENSDARG00000062198). We used the isoform pcm1-201 (ENSEMBL ENSDART00000149026). Both N- and C-terminal tagged versions of Pcm1 were generated and tested. GFP-linker-Pcm1, when expressed in the embryos, caused embryonic lethality, suggesting that N-terminal tagged Pcm1 dominantly interfered with embryonic development. The C-terminal tagged Pcm1, which did not interfere with embryogenesis, was used in our study. mRNAs encoding Pcm1-linker-GFP were microinjected into one blastomere of 16-cell stage embryos to achieve sparse labeling. At ∼22 hpf, an antibody against the Notch ligand Dld was injected into embryonic brain ventricles to label internalized endogenous Dld as previously described ^29^ (**Fig. 1g**). Pcm1 expression level varied; in most RGPs, the Pcm1-GFP reporter formed puncta (of variable sizes depending on expression levels). These puncta were in proximity to Dld endosomes and together they moved toward the posterior side (**Fig. 1h, 1i**, and **Supplementary Videos 1-3**). By telophase, most Pcm1-GFP and Dld endosomes were enriched in the posterior daughter (**Fig. 1j**). Together, these results uncover a previously unknown asymmetry of Pcm1 at the centrosomes and its localization in the central zone near Dld endosomes followed by enrichment in the posterior daughter.

### Requirement of *pcm1* for asymmetric distribution of endosomes in mitotic RGPs

In zebrafish, asymmetric segregation of Dld endosomes in mitotic RGPs is dependent on the polarity regulator Par-3 and the dynein motor complex ^12^. To understand how Pcm1 might contribute to endosome asymmetry, we labeled the internalized Dld as schematized in **Fig. 1g** and performed *in vivo* time-lapse imaging in control and *pcm1*-deficient embryos. A significant reduction of dividing RGPs was observed in both *pcm1* MOs and KOs. Intriguingly, perpendicular divisions were reduced with a corresponding increase of non-perpendicular divisions that are often associated with differentiation (**Extended Data Fig. 2a, 2c**). In the *pcm1*-deficient RGPs that were able to divide, reduced numbers of Dld-containing endosomes were observed (**Extended Data Fig. 2b**), suggesting a role of Pcm1 in facilitating endocytosis.

Despite a reduction of Dld endosomes in *pcm1*-deficient RGPs, enough were present for characterizing their intracellular dynamics. In most WT mitotic RGPs, internalized Dld first converged toward the central zone, then moved in a polarized direction toward the posterior side. However, in *pcm1*-deficient RGPs, asymmetric segregation of Dld endosomes into the posterior daughter was significantly reduced (**Fig. 2a, b**, and **Supplementary Videos 4-6**). Analysis of single-endosome dynamics showed that endosome trajectories were not posterior-bound in *pcm1*-deficient mitotic RGPs (**Fig. 2c**). Their distribution shifted significantly anteriorly (**Fig. 2d**). Moreover, the average velocity of *pcm1*-deficient endosomes was significantly higher than controls (**Fig. 2e**), suggesting that the deficiency in posterior-bound movement is due to a lack of directionality, not motility of endosomes. Together, these results indicate that Pcm1 is required to regulate the polarized dynamics of Dld endosomes toward the posterior. Additionally, Pcm1 plays a role in mitotic cell cycle entry and endocytosis.

**Fig 2.**
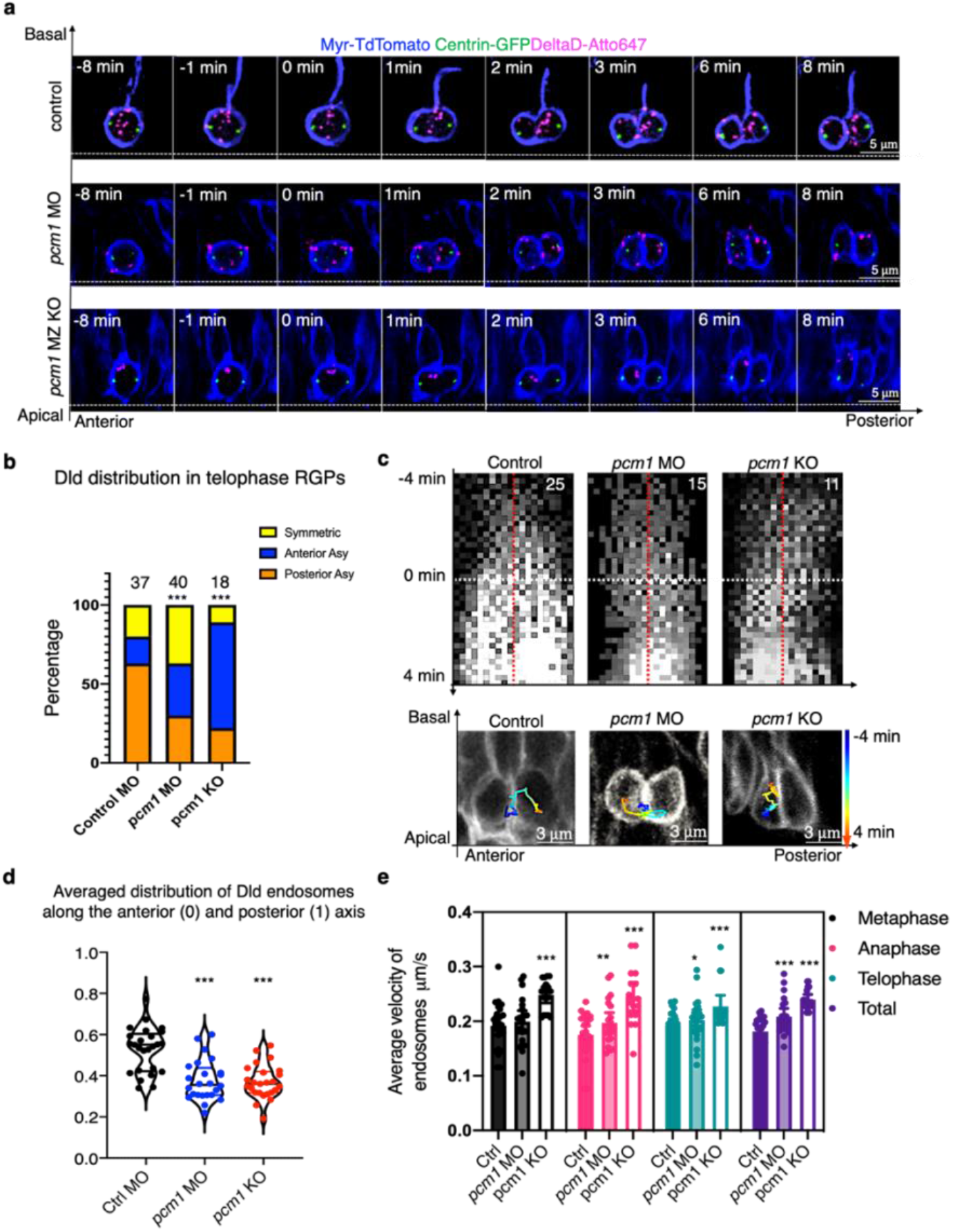
*In vivo* time-lapse imaging reveals the requirement of Pcm1 for Dld endosome movement toward the posterior in zebrafish embryonic forebrain mitotic RGPs. **a**. Time-lapse imaging montages in control MO, *pcm1* MO, and *pcm1* KO. Membrane is marked with Myr-TdTomato (pseudo-colored blue), centrosomes are marked with Centrin-GFP (pseudo-colored green), and internalized Dld endosomes are marked with Dld-atto647 (pseudo-colored magenta). T= 2 min denotes the onset of telophase (determined by the membrane abscission between the daughter cells). Each fame is the MIP of 5 confocal z-stacks (1 μm z-step). White dashed lines indicate the apical-basal and anterior-posterior axes. **b**. Statistics of the distribution of internalized Dld in telophase RGPs. *** *p* < 0.0001, χ^2^ test (chi-square = 73.46, df = 4). **c**. Kymographs (top) and representative trajectories (bottom) of Dld endosomes, tracked from -4 min to 4 min (the starting point of anaphase is 0 min, indicated by white dashed line). The red dashed lines on the kymograph images indicate the center of the cell registered by centrosomes. Compared to controls, both posterior enrichment and posterior-directed movements of Dld endosomes were reduced in *pcm1* MO and *pcm1* KO. **d**. Averaged distribution of Dld endosomes along the A-P axis; each dot representing the value from one RGP at telophase (T = 4 min). The positions of centrosomes are used for cell registration. Compared to controls, the distribution of Dld endosomes in both *pcm1* MO and *pcm1* KO RGPs was significantly more anterior. *** *p* < 0.001, n = 25 telophase RGPs per group, unpaired two tailed t-test. **e**. Average velocity of Dld endosomes in dividing RGPs (T = -4 min to T = 4 min). Significantly higher velocity was detected in *pcm1* MO and KO RGPs compared to control. The T frame interval is 12 sec. Each dot represents the average velocity of all Dld endosomes in a single RGP. *** p < 0.001; ** p < 0.01, * p < 0.05; Control group, n=25 RGPs; pcm1 MO, n=21 RGPs; pcm1 KO, n = 17 RGPs.

### Requirement of *pcm1* for regulating progenitor division mode and daughter cell fate revealed at clonal resolution

To determine the impact of Pcm1 on RGP division mode and daughter cell fate, we used a transgenic line *Tg[HuC-GFP;Ef1a-H2B-mRFP]*, which marks neurons green and all nuclei red (**Fig. 3a**). Quantification of neuron numbers relative to total nuclei in the telencephalon showed a significant relative increase of neurons in *pcm1*-deficient embryos (**Fig. 3b**), although the overall number of nuclei appeared decreased (**Fig. 3a**), consistent with reduced proliferation observed in *pcm1*-deficient embryos.

To determine whether the relative increase of neuronal production in *pcm1*-deficient embryos is due to defects in the asymmetry of Dld endosomes resulting in altered modes of cell division, we performed clonal analysis of RGP division. One-cell stage *Tg[HuC-GFP]* embryos were injected with either control or *pcm1* MO, followed by H2B-mRFP mRNA injection into one blastomere at the 16-32-cell stage. *In vivo* long-term time-lapse imaging was carried out to track mitotic RGPs for ∼10 hours (**Fig. 3c**). Three types of divisions were observed (**Fig. 3d-e, and Supplementary Videos 7-9**): 1) progenitor/progenitor (P/P), where both daughters remained HuC-GFP^-^ 10 hr after division; 2) progenitor/neuron (P/N), where one daughter remained a progenitor and the other became a neuron; 3) neuron/neuron (N/N), where both daughters became HuC-GFP^+^. In control MO-injected embryos, 42% of RGP divisions were P/P, 39% P/N, and 19% N/N (n=72). In *pcm1* MO embryos, there was a decrease of P/P and P/N divisions and a corresponding increase of N/N divisions, which now accounted for 49% of observed divisions (n=45) (**Fig. 3d-e**).

**Fig 3.**
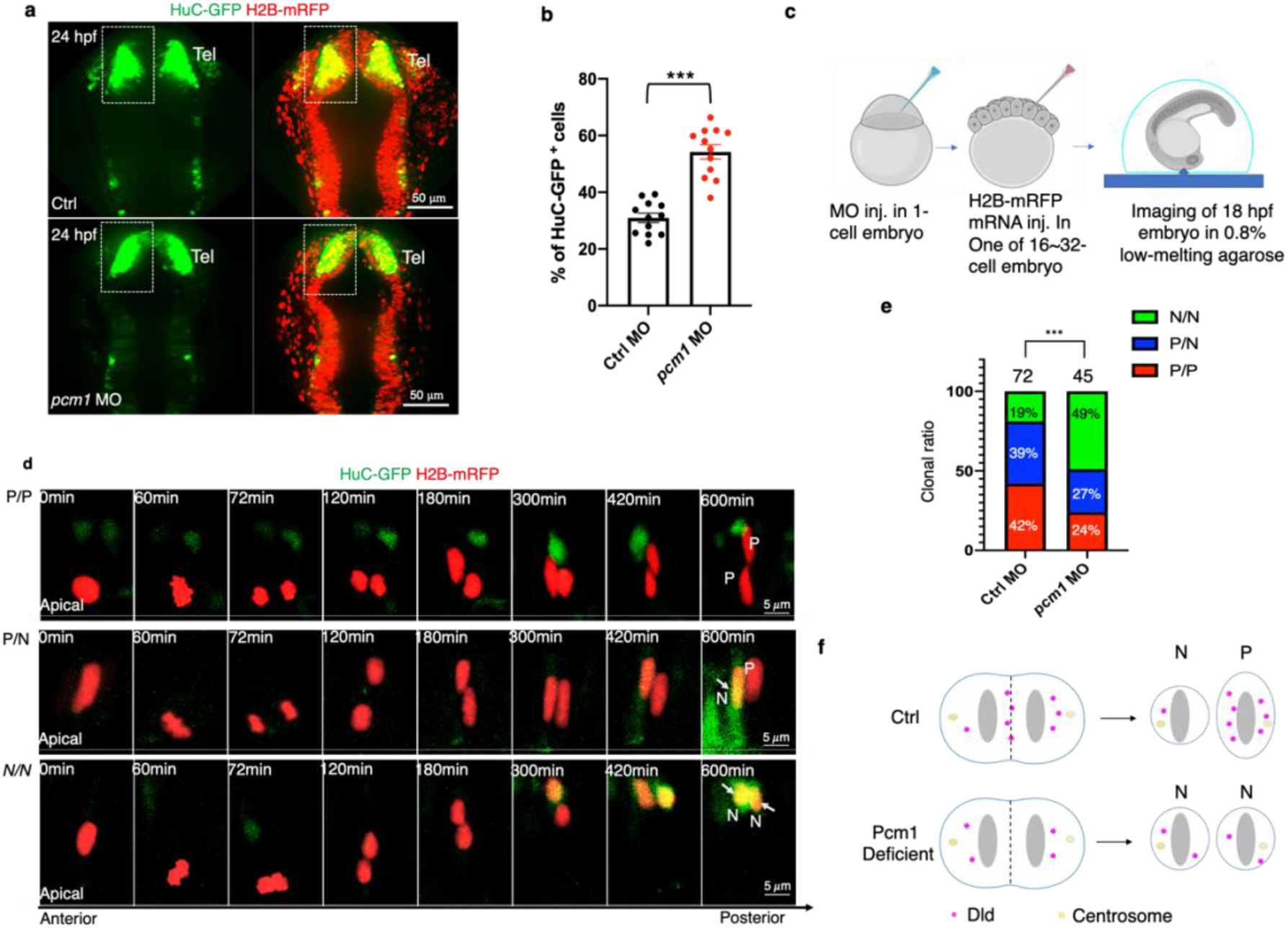
Pcm1 is required to maintain proliferative RGPs in the developing zebrafish forebrain. **a**. Live imaging of HuC-GFP^+^ cells of 24 hpf *Tg[HuC-GFP: H2B-mRFP]* embryos. Differentiating neurons in the telencephalon (Tel) are boxed in both control MO and *pcm1* MO embryos. Each image is the MIP of 10 z-planes. The z-step is 1 μm. Scale bars: 50 μm. **b**. Statistics show a significant increase of HuC-GFP^+^ cells relative to the total telencephalic nuclei (H2B-mRFP^+^) in the boxed area of (a). n=12, ***, p < 0.001. **c**. Schematic of sparse labeling and time-lapse imaging. **d**. Montages of time-lapse sequence of images showing clonally labeled RGPs giving rise to progenitor (P) daughter and/or neuron-like (N) daughter cells. Each image is the MIP of 10 z-planes. The z-step is 1 μm. The time interval between each volume of z-stacks is 6 min and the total acquisition time is ∼10 hours. **e**. Statistics show that RGPs in *pcm1* MO embryos underwent more N/N and fewer P/P and P/N divisions. *** p < 0.001, χ^2^ test (Chi-square = 19.46, df = 2). **f**, schematic summarizing both the Dld endosome (Fig. 2) and cell fate phenotypes (this Figure).

Additionally, we performed EdU pulse chase experiments in *pcm1* KO embryos (**Extended Data Fig. 3a**). Consistent with reduced number of mitotic RGPs (**Extended Data Fig. 2a**), we found decreased EdU labeling, indicative of fewer cells entering S-phase in *pcm1*-deficient embryos (**Extended Data Fig. 3b, 3d**). After 24-hr chase, the proportion of EdU^+^Hu^+^ in all EdU^+^ cells was significantly increased in both the forebrain and spinal cord, suggesting an increased neuronal production at the expense of progenitors in *pcm1*-deficient embryos (**Extended Data Fig. 3c, 3e**). Overproduction of neurons at the expense of progenitors has also been observed in *pcm1*-deficient mice ^30^. Together, findings in zebrafish reveal at an unprecedented clonal resolution that Pcm1 maintains RGP progenitor fate by promoting P/P and P/N division and/or suppressing the N/N division. This is likely due to the requirement of Pcm1 for the asymmetric segregation of Dld endosomes (as shown in **Fig. 2**): In the absence of Pcm1, Notch signaling is defective, thereby resulting in a decrease of P/P and an increase of N/N divisions.

### Co-localization of Pcm1 with Par-3, Rab5b, and Rab11a on Dld endosomes revealed via high resolution expansion microscopy

To understand how Pcm1 directs Dld endosome movements toward the posterior side, we analyzed its sub-cellular co-localization with other proteins in mitotic RGPs, first by conventional immunofluorescent microscopy (**Extended Data Fig. 4**), then high resolution expansion microscopy (**Fig. 4, 5**). In developing zebrafish forebrain mitotic RGPs at anaphase, Pcm1 was detected in the central zone, where the polarity regulator Par-3, Dld endosomes, the dynein complex component Dlic1, the dynactin subunit P150, and the recycling endosome markers Rab11a and the early endosome marker Rab5b were also detected. The presence of Rab5b in the central zone was much less compared to Rab11a, suggesting that the central zone mostly harbors recycling endosomes (**Extended Data Fig. 4a-b**). Quantification of co-localization coefficient ^31^ among pairs of proteins uncovered co-localization of Pcm1 with many of these proteins, in both the central zone and the surrounding cytoplasm (**Extended Data Fig. 4c**). Both Pcm1 and Par-3 showed more co-localization with Rab11a in the central zone and with Rab5b in the surrounding cytoplasm. *In vivo* co-immunoprecipitation further showed that Pcm1 formed complexes with Par-3, Dlic1, and Dld, Rab11a, and Rab5b (**Extended Data Fig. 5**).

To visualize protein co-localization at a higher resolution, we performed label retention expansion microscopy (LR-ExM). This technology enables us to detect cytoplasmic Par-3 that decorate Dld endosomes ^12^. Endosomes were visualized in five z-plane projected images (**Fig. 4a, right**). Intriguingly, Pcm1 intercalated with Par-3 on Dld endosomes (**Fig. 4a, and Extended Data Fig. 6a**). Dld endosomes decorated with both Par-3 and Pcm1 were significantly more in the central zone than in the surrounding cytoplasm (**Fig. 4b**).

**Fig 4.**
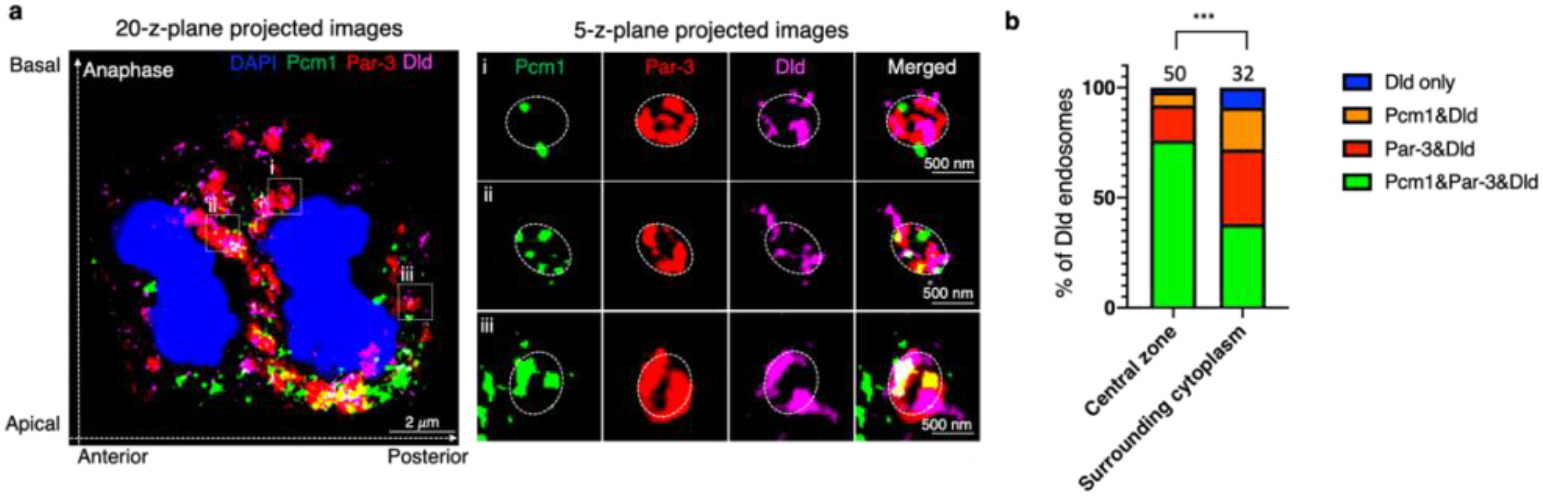
Label retention expansion microscopy (LR-ExM) uncovers Pcm1 colocalization with Par-3 on Dld endosomes in zebrafish embryonic forebrain mitotic RGPs. **a**. Anaphase RGP immuno-stained with anti-Par-3, anti-Pcm1, anti-Dld and DAPI from 28 hpf embryo cryosections. The left panel shows whole cell image (MIP of 20 z-planes), and the right panel shows enlarged views (i, ii and iii) of endosomes (MIP of 5 z-planes). The z-step is 0.26 μm. Scale bars denote the biological size. **b**. Statistics of the percentage of Par-3, Pcm1 and Dld-containing endosomes. All endosomes (50 at the central zone and 32 from the surrounding cytoplasm) are from 18 RGPs. *** *p* < 0.0001, χ ^2^ test (Chi-square = 30.36, df = 3).

Using Rab5b and Rab11a to visualize the early and recycling Dld endosomes respectively, we detected an association of Pcm1 with both markers on Dld endosomes (**Fig. 5a-b, Extended Data Fig. 6b-c**). P150, a subunit of the dynactin complex, was also found on Dld endosomes (**Extended Data Fig. 6c**). Quantifications uncovered that Dld endosomes in the central zone were enriched with Pcm1 and Rab11a, whereas Dld endosomes in the surrounding cytoplasm were enriched with Pcm1 and Rab5b. Together, these findings reveal at high resolution that Pcm1 is associated with Dld recycling endosomes that are decorated with Par-3 and dynein/dynactin complex in the central zone.

**Fig 5.**
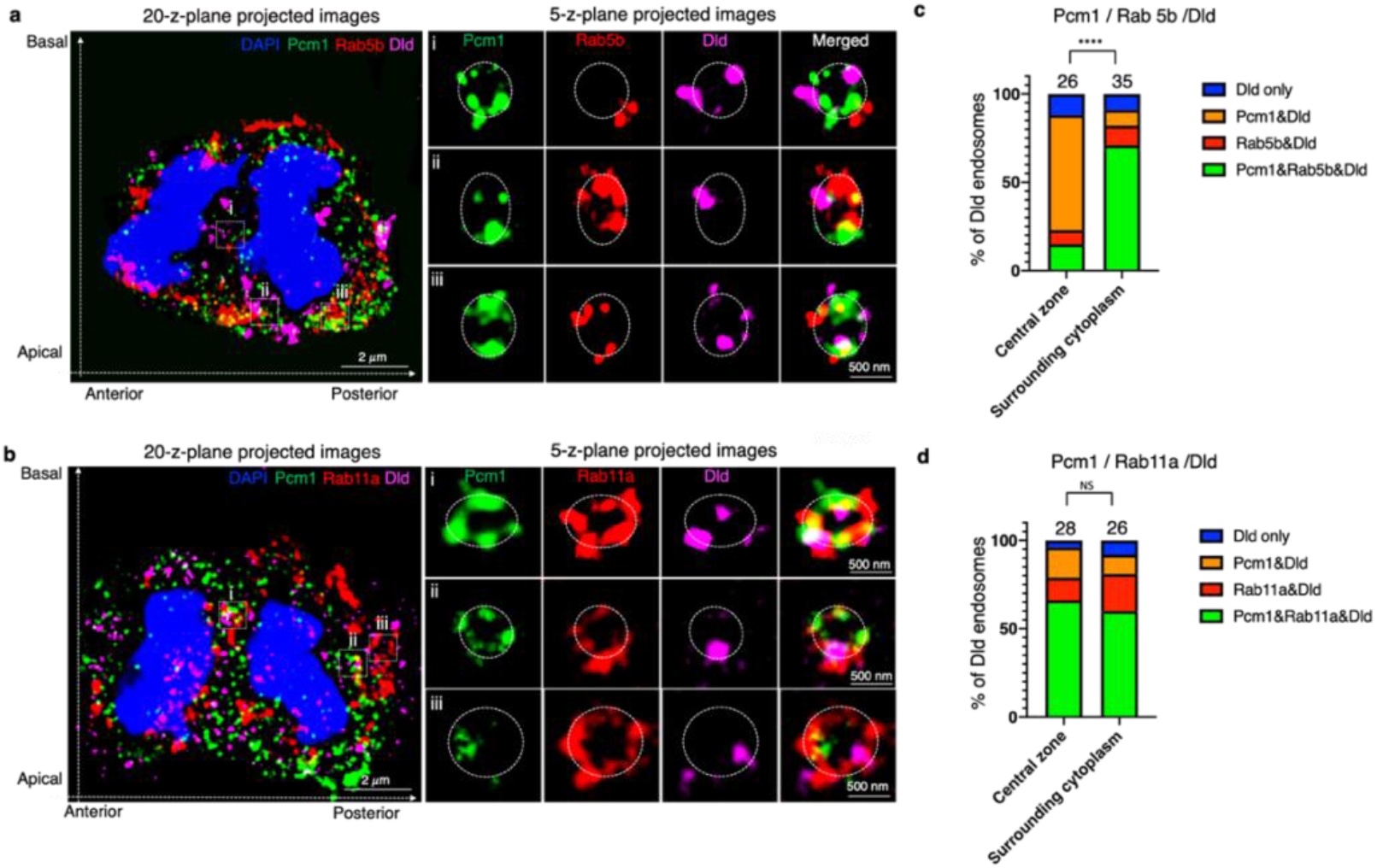
LR-ExM uncovers Pcm1 colocalization with Rab5b and Rab11a on Dld endosomes in zebrafish embryonic forebrain mitotic RGPs. **a-b.** Anaphase RGP immuno-stained with anti-Pcm1, anti-Rab5b (**a**) or anti-Rab11a (**b**), anti-Dld, and DAPI from 28 hpf embryonic forebrain cryosections. The left panels show whole cell image (MIP of 20 z-planes); the right panels show enlarged views of endosomes (i, ii and iii) (MIP of 5 z-planes). The z-step is 0.26 μm. Scale bars denote the biological size. **c**. Statistics of the percentage of Rab5b, Pcm1 and Dld-containing endosomes identified in anaphase RGPs. All endosomes (26 in the central zone and 35 from the surrounding cytoplasm) are from 16 RGPs. ** *p* <0.0001, χ^2^ test (Chi-square = 79.75, df = 3). **d**. Statistics of the percentage of Rab11a, Pcm1 and Dld-containing endosomes identified in anaphase RGPs. All endosomes (28 in the central zone and 26 from the surrounding cytoplasm) are cfrom 15 RGPs. NS, *p* = 0.1881, χ^2^ test (Chi-square = 4.787, df = 3).

### Requirement of Pcm1 for the formation of Par-3-Dynein macromolecular complexes on Dld endosomes

To determine what role Pcm1 might play on Dld endosomes, we performed high-resolution expansion microscopy and analyzed anaphase/telophase RGPs in *pcm1*-deficient embryos. In control, most Dld endosomes were decorated with both Par-3 and the dynein complex component Dlic1. In *pcm1*-deficient embryos, however, we observed a significant decrease of Par-3^+^Dlic1^+^ Dld endosomes and a corresponding increase of Par-3^+^-only and Dlic1^+^-only Dld endosomes (**Fig. 6a-b**). Moreover, the amount of Dld in these Par-3^+^-only and Dlic1^+^-only Dld endosomes was significantly less than that in Par-3^+^Dlic1^+^ Dld endosomes (**Fig. 6c**). These findings suggest that Pcm1 is required to bring together both Par-3 and Dlic1 on Dld endosomes; in its absence, most Dld endosomes have either Par-3 or Dlic1 but not both. The amount of Dld is further reduced in these Pcm1-deficient Par-3-only or Dlic1-only endosomes. These findings suggest that Pcm1 is required for macromolecular complex formation on Dld endosomes and for the integrity of these endosomes.

**Fig 6.**
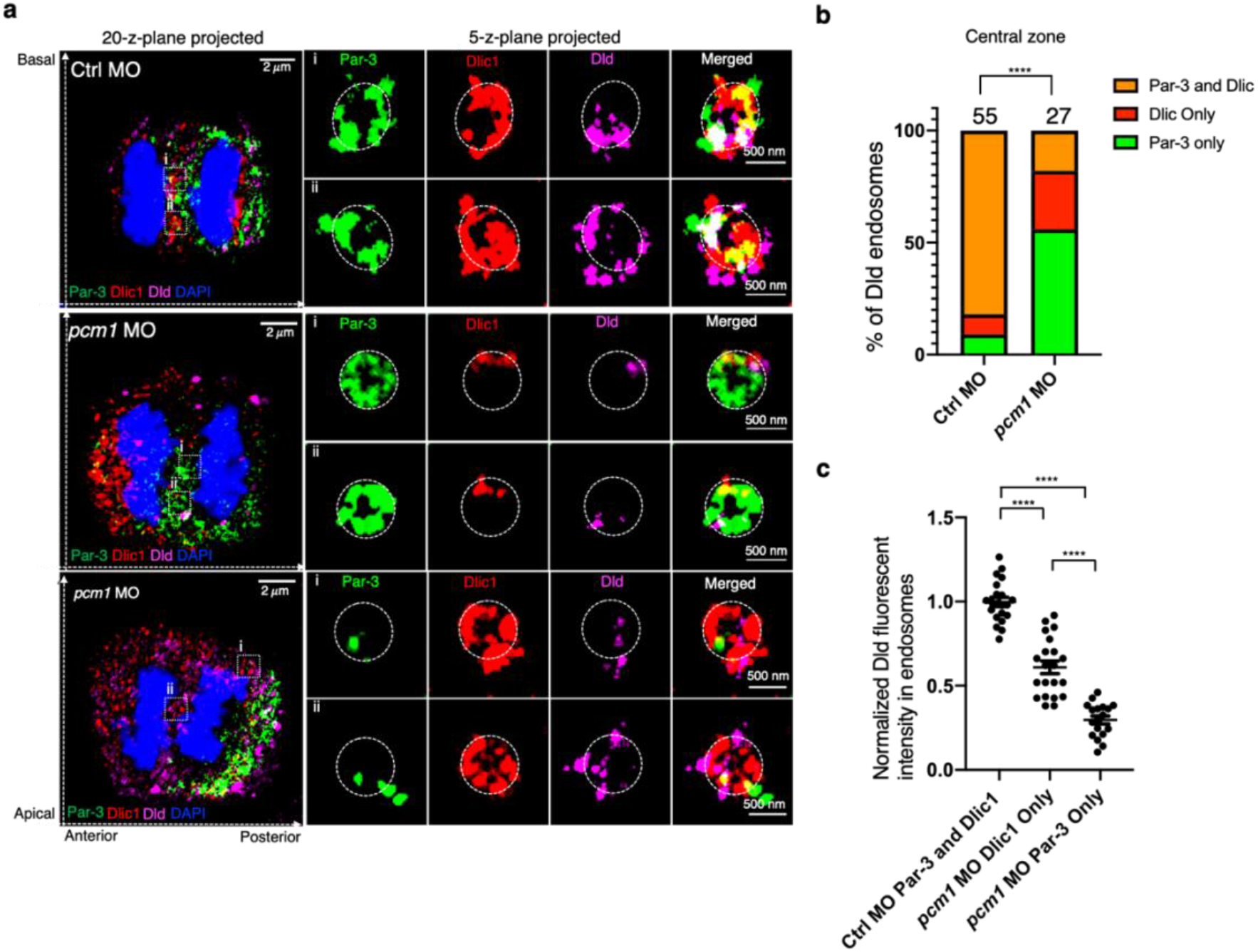
Pcm1 is required for the colocalization of Par-3 and Dlic1 on Dld endosomes in zebrafish embryonic forebrain mitotic RGPs. **a**. LR-ExM shows Par-3 and Dlic1 on Dld endosomes. RGPs from control MO (upper panel) and *pcm1* MO (lower two panels) are shown. The left panels show whole cell images (MIP of 20 z-planes); the right panels show enlarged views (i and ii) (MIP of 5 z-planes). The z-step is 0.26 μm. The dashed lines with arrows denote anterior-posterior axis and apical-basal axis. Scale bars denote the biological size. **b**. Statistics of the percentage of Dld endosomes with Par-3, Dlic1, or both. Endosomes were analyzed from 6 RGPs of three independent experiments for each group. **** *p* < 0.0001, χ^2^ test (Chi-square = 83.20, df = 2). **c**. Quantification of Dld fluorescent intensity on endosomes. The anti-Dld immunofluorescent intensity was first normalized to that of DAPI, and then to that of Par-3^+^Dlic1^+^ Dld endosomes from the control MO. All endosomes are from six RGPs for each group, collected from three independent experiments. Unpaired t-test, **** *p* < 0.0001, n =21 endosomes for each group.

### Requirement of Pcm1 for the trafficking from early to recycling endosomes

Given Pcm1’s association with general endosomal proteins including Rab11a and Rab5b, we reasoned that its role on endosomes might not be limited to those containing Dld. To test this, we analyzed endosomes marked with Rab11a or Rab5b in anaphase/telophase mitotic RGPs in *pcm1*-deficient embryos. In control RGPs triple immuno-stained with Par-3, Rab11a, and P150 antibodies, most endosomes in the central zone had Par-3, Rab11a, and P150. However, in *pcm1*-deficient embryos, Par-3^+^Rab11^+^P150^+^ endosomes were significantly reduced, with a corresponding increase of Rab11a-P150 and Par-3-P150 co-labeling (**Fig. 7a-b**). These observations suggest that Pcm1 is required to bring together Par-3 and P150 on recycling endosomes.

Rab5^+^ endosomes colocalizing with Par-3 and p150 were less frequently observed in the central zone in control anaphase/telophase mitotic RGPs. However, the number of Rab5^+^Par-3^+^P150^+^ endosomes significantly increased in the central zone of *pcm1*-deficient anaphase/telophase mitotic RGPs (**Fig. 7c-d**). These data suggest that Pcm1 is required for the trafficking from Rab5^+^ early endosomes to Rab11^+^ recycling endosomes; this deficit possibly contributes to the decreased Rab11^+^Par-3^+^P150^+^ endosomes in the central zone of *pcm1*-deficient mitotic RGPs.

### Dysregulation of genes involved in endocytosis and cell differentiation upon Pcm1 disruption

Consistent with the localization of Pcm1 to endosomes and its function in regulating endosome dynamics, we detected changes of gene expression related to endocytosis and cellular differentiation at the transcriptomic levels upon Pcm1 disruption. RNA-seq was performed on control and *pcm1* KO embryos at ∼ 24 hpf. Among the 1,176 genes with decreased and 1,034 genes with increased expression in the *pcm1* KO embryos, gene ontology (GO) analysis uncovered a significant enrichment for the genes involved in endocytosis and vesicle-mediated transport (**Extended Data Fig. 7a**). They include genes encoding proteins involved in clathrin coated vesicle formation and endocytosis (e.g., *caly, ston2, ap1m2, ap3b1a*), receptor-mediated endocytosis (e.g., *lmbr1l, dbnla*), vesicle trafficking (e.g., *stxbp5l, dop1a, ap2m1b, flot1a, sytl1*), exocytosis (e.g., *cplx2l, stxbp5l, anaxa1d, syt7b*), endosomal sorting (e.g., snx5, snx2, *ston2, ap1m2*), or localized to endosomes (e.g., *stard3, rab5b, vps8*) (**Extended Data Fig. 7b-c**). Together, these results support a role of Pcm1 in regulating endocytosis and endosome dynamics, which is critical for proper progenitor fate and cellular differentiation.

**Fig 7.**
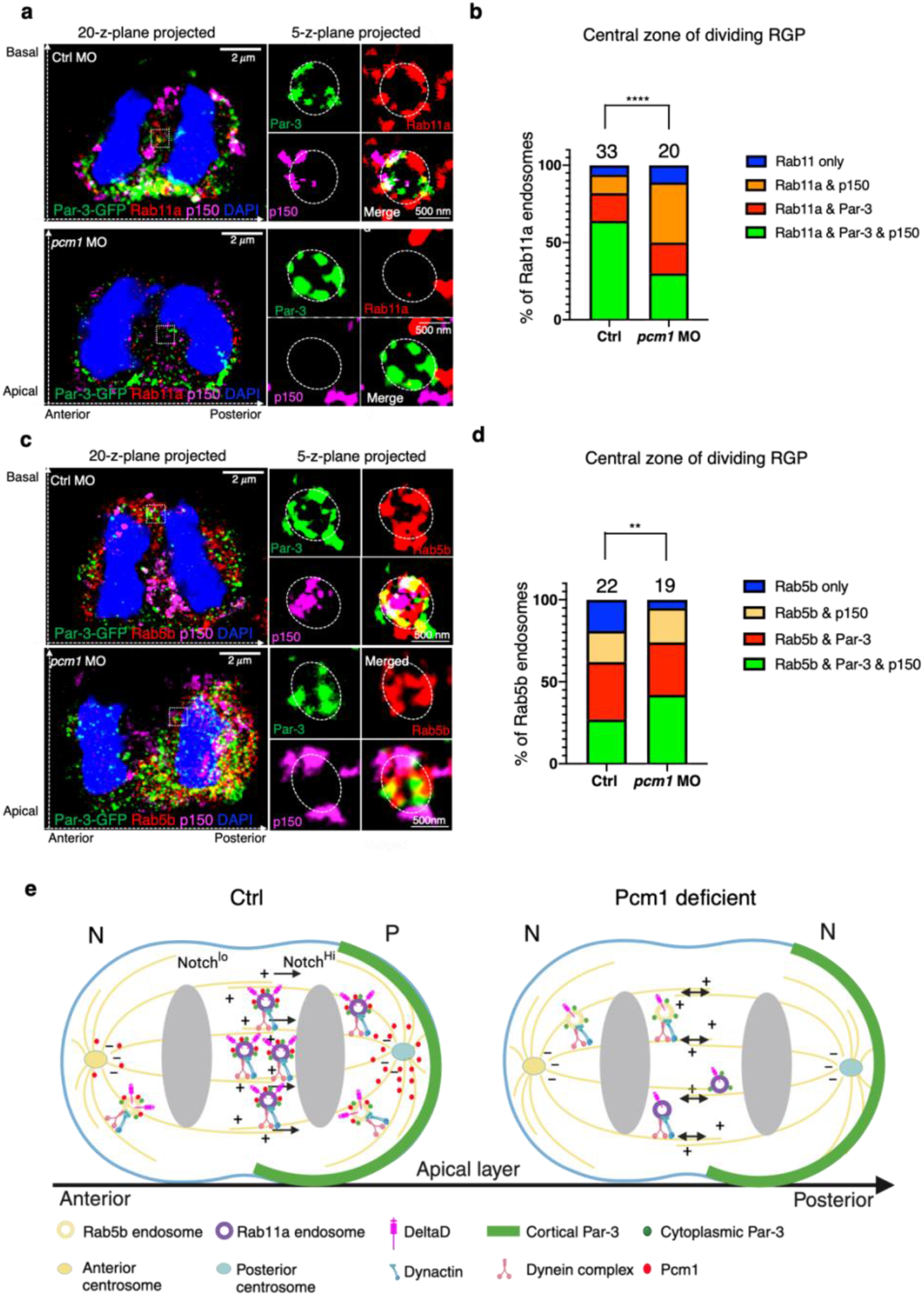
Pcm1 is required for trafficking from Rab5 early endosomes to Rab11 recycling endosomes in zebrafish embryonic forebrain mitotic RGPs. **a**. LR-ExM shows Par-3 with Rab11a and P150 on recucling endosomes. **b**. Statistics of endosomes with different compositions of Par-3, Rab11a, and/or P150 in the central zone of control and *pcm1*-deficient RGPs. Endosomes were from six RGPs for each group, collected from three independent experiments. **** *p* < 0.0001, χ^2^ test (Chi-square = 28.17, df = 3). **c**. LR-ExM shows Par-3 with Rab5b and P150 on early endosomes. **d**. Statistics of endosomes with different compositions of Par-3, Rab5b and/or P150 in the central zone of control and *pcm1*-deficient RGPs. Endosomes were from six RGPs for each group, collected from three independent experiments. ** p = 0.0086, χ^2^ test (Chi-square = 11.66, df = 3). For both **a** and **c**, the left panels show whole cell images (MIP of 20 z-planes); the right panels show enlarged views of endosomes (MIP of 5 z-planes). The z-step is 0.26 μm. Scale bars denote the biological size. **e**, Schematic summary of Pcm1’s role in macromolecular complex formation and dynamics of Dld endosomes and progenitor fate. N: neuron; P: progenitor.

### Asymmetry and colocalization of PCM1 with PARD3 and the recycling endosome component RAB11A in hiPSC-derived neural progenitors

To determine whether the association of PCM1 with endosomes might be a conserved phenomenon, we examined the distribution of PCM1 in hiPSC-derived neural progenitors. Two well-characterized hiPSC lines, WTC11 ^32^ and KOLF2.1J ^33^, were used for differentiation toward 2D neural rosettes and 3D organoids with dorsal forebrain characteristics, using a previously published method ^34^. The 3D organoids were obtained after 25 days in culture and the 2D neural rosettes were harvested for analysis after 21 days in culture (**Fig. 8a**). We analyzed over 20 neural rosettes and 8 organoids from each line. This was repeated in four independent differentiation experiments for each line.

**Fig 8.**
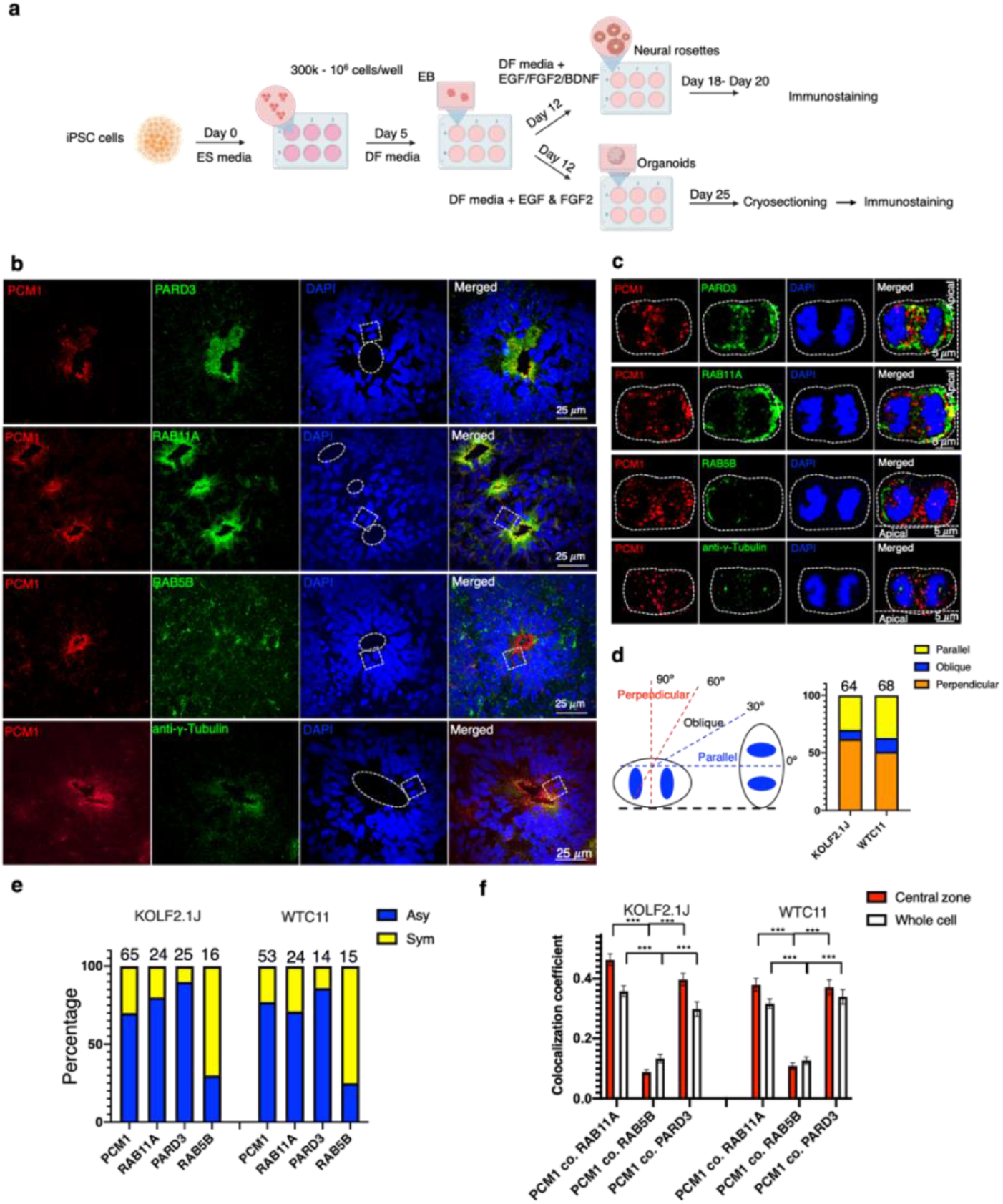
PCM1 asymmetry and colocalization with PARD3 and RAB11A in mitotic neural progenitors of human brain organoids. **a**. Schematic showing the derivation of forebrain organoids and neural rosettes from 2 hiPSC lines, KOLF2.1J and WTC11. **b**. Detection of PCM1, PARD3, RAB11A, and RAB5B in mitotic neural progenitor cells (NPCs) of forebrain organoid cryosections (14 μ m thickness). The ventricular zone (VZ)-like regions are marked with dashed circles. MIP of 20 z-planes is shown. Z-step is 0.26 μm. Scale bar, 25 μ m. **c**. Dividing NPCs (marked with dashed rectangles in b). MIP of 10 z-planes is shown. Z-step is 0.26 μ m. Scale bar, 5 μm. White dashed lines in the right panel demarcate the VZ surface. **d**. Statistics of division orientation of NPCs. The cartoon represents three main types of division orientations according to the angles between the cleavage plane and the VZ apical surface. **e**. Statistics of PCM1, RAB11A, PARD3 and RAB5B distribution in anaphase/telophase NPCs. In forebrain organoids derived from both hiPSC lines, PCM1, RAB11A, and PARD3 were asymmetrically distributed in most NPCs. In contrast, RAB5B was symmetrically distributed in most NPCs. **f**. Statistics of colocalization coefficients of PCM1 with PARD3, RAB11A and RAB5B in the anaphase/telophase NPCs of forebrain organoids. In the central zone, colocalization coefficients of PCM1 with RAB11A (PCM1 co. RAB11A) and PCM1 with PARD3 (PCM1 co. PARD3) are significantly higher than PCM1 with RAB5B (PCM1 co. RAB5B). *** *p* < 0.001, unpaired t test. For KOLF2.1J NPCs, n= 24 for PCM1 co. RAB11A.; n=25 for PCM1 co. PARD3; n=16 for PCM1 co. RAB5B. For WTC11 NPCs, n= 24 for PCM1 co. RAB11A.; n=14 for PCM1 co. PARD3; n=15 for PCM1 co. RAB5B.

The organoids consisted of multiple pseudo-stratified ventricular zone (VZ)-like progenitor regions and expressed dorsal telencephalic progenitor markers PAX6, the neural stem/progenitor marker NESTIN, intermediate progenitor cell (IPC) marker TBR2, and nascent neuronal marker HUC (**Extended Data Fig. 8a**). The neural rosettes expressed PAX6, NESTIN and HUC (**Extended Data Fig. 8b**). We also validated our custom chicken anti-PCM1 antibody in neural rosettes (**Extended Data Fig. 8c**), which showed identical staining patterns as a previously published rabbit anti-PCM1 antibody ^21^.

In cryo-sectioned brain organoids, PCM1, PARD3, and RAB11A were enriched and co-localized in the VZ-like progenitor regions, whereas only a partial overlap was observed for PCM1, RAB5B, and γ-tubulin (a centrosome marker) (**Fig. 8b**). A close examination of mitotic neural progenitors at late anaphase or telophase found that most displayed asymmetric distribution of PARD3, PCM1, and RAB11A, but not RAB5B (**Fig. 8c, 8d**). Similar to previous findings ^35, 36^, we also noted an increased percentage of progenitors that had their cleavage planes parallel to the VZ surface (**Fig. 8e**), compared to divisions of zebrafish and mouse embryonic neural progenitors with cleavage planes mostly perpendicular to the VZ. Quantification of protein co-localization showed the co-localization of PCM1 with PARD3 and RAB11A, but much less with RAB5B, in both the central zone and the whole cell (**Fig. 8f**). Similar observations were made in the hiPSC-derived neural rosettes (**Extended Data Fig. 9**). Together, these data suggest that the association of PCM1 with PARD3 and the recycling endosome protein RAB11A and their asymmetric distribution in mitotic neural progenitors are conserved from fish to humans.

## Discussion

In this study, we have uncovered an asymmetric distribution of Pcm1 at centrosomes and its role in polarizing endosome dynamics during ACD of RGPs. At metaphase, Pcm1 is detected as an amorphous pericentriolar cloud enriched at the posterior centrosome. At anaphase, Pcm1 is localized to the Dld endosomes and is required for their enrichment in the posterior self-renewing daughter. Pcm1 is necessary for the formation of macromolecular complexes composed of Par-3 and dynein on Dld endosomes. This molecular action likely underscores the requirement of Pcm1 in polarized endosome dynamics and in the maintenance of progenitor fate. We further show that PCM1 co-localizes with PARD3 and the recycling endosome marker RAB11A and is asymmetrically distributed in most hiPSC-derived mitotic neural progenitors, suggesting a possibly conserved role of PCM1 in regulating endosome dynamics in human neural progenitors. Previous studies have found recycling endosomes in the pericentrosomal area ^37^, intimately contacting the appendages of the mother centriole ^38^. These observations provide a structural basis for the coordinated asymmetry between centrosomes and endosomes during ACD, which was discovered for the first time to our knowledge in this study. Our findings suggest that Pcm1 plays a key role in coordinating multi-faceted intracellular asymmetry, from the cortical domains to the centrosomes, and to the endosomes, contributing to the maintenance of RGP progenitor fate (**Fig. 7e**). While it is comprehensible that asymmetry of Notch signaling can confer distinct daughter cell fate, it has been unclear how the asymmetric inheritance of centrosomes might impact cell fate. Our finding of a crosstalk between centrosomes and endosomes, mediated by Pcm1 during ACD, suggest a new mechanism, by which the asymmetry at the centrosomes, which could be involved in regulating the polarized endosome dynamics and the asymmetry of Notch signaling, lead to distinct daughter cell fate.

In the developing zebrafish forebrain, mitotic RGPs inherit an apical-basal polarity from their precursor neuroepithelial progenitors, with Par-3 localized to the apical domain. However, the mitotic spindle in most RGPs is positioned such that the cleavage plane bisects the apical domain. This division orientation necessitates a new axis of asymmetry in mitotic RGPs, with cortical Par-3, centrosome Pcm1, and Dld endosomes all polarized along the A-P axis. Furthermore, Pcm1, Par-3, and dynein components, are all directly associated with Dld endosomes, where Pcm1 facilitates the trafficking from early to recycling endosomes and promotes co-localization of Par-3 and dynein. What extrinsic signals polarize RGPs along the A-P axis and what additional factors might help transduce such signals among different cellular compartments, are interesting questions for future investigations.

Previous studies in the developing mammalian cortex ^5, 14, 39^ and our findings in hiPSC-derived brain organoids suggest a considerable degree of conservation in the behavior of vertebrate RGPs. Further comparative analyses will provide deep insights into the development and evolution of this remarkable cell type.

## Methods

### Zebrafish strains and maintenance

Wild-type embryos were obtained from natural spawning of AB adults maintained according to established protocols ^40^. Embryos were raised at 28.5 °C in 0.3 x Danieau’s embryo medium (30x Danieau’s embryo medium contains 1740 mM NaCl, 21 mM KCl, 12 mM MgSO4•7H2O, 18 mM Ca(NO_3_)_2_, 150 mM HEPES buffer). Embryonic ages were described as hours post-fertilization (hpf). To prevent pigment formation, 0.003% Phenylthiourea (PTU) was added into the medium cultured with 24 hpf embryos. The following zebrafish mutants and transgenic lines were used: Tg [ef1α:Myr-Tdtomato] and Tg [HuC:GFP] ^6, 12^. All animal experiments were approved by the Institutional Animal Care and Use Committee (IACUC) at the University of California, San Francisco, USA.

### Generation of *pcm1* knockout zebrafish

nls-zCas9-nls RNA was *in vitro* transcribed from a plasmid (a gift from Wenbiao Chen) ^41^, and injected with two sgRNAs (*TCC-*ATTCACTTAGAGACCAGACCC, Exon 8; *CCT-* CCAATAATAGAGATGGCCGC, Exon 11) into one-cell stage zebrafish embryos. The PCR primers for genotyping are 5’-TTG ACT CGC CTG TAA CTT GTT G-3’ and 5’-CAA TGA GGT TAG TGT GGA ATC C-3’, which flanked the regions from Exon 8 to Exon 15 of the zebrafish *pcm1* gene. The sequencing primers used are 5’-GGA AGC TGA AGG AGG TGC ACA A 3’ and 5’-TGG TGC AGG TGA TAT TCT AGT CA-3. The F2 generations of homozygous male and female *pcm1* knockout adult fish were used to generate the maternal zygotic *pcm1* KO.

### Primary antibodies

Rabbit anti-γ-tubulin [Sigma Cat# T5192, Research Resource Identifier (RRID): AB_261690, 1:500 for immunostaining]. Rab5b antibody (rabbit polyclonal, Invitrogen Cat# PA5-44574, RRID: AB_2608403, 1:500 for immunostaining), Rab11a (rabbit polyclonal, Thermofisher PO# 715300, RRID: AB_2533987, 1:500 for immunostaining), Anti-HuC/D (mouse monoclonal, Thermofisher PO# A-21271, RRID: AB_221448, 1:500 for immunostaining), Mouse anti-Dld (Abcam, ab73331; RRID, catalog number: AB_1268496; lot GR115501-3, 1:200 dilution for immunostaining); Chicken anti-GFP (Abcam, catalog number: ab13970; RRID:AB_300798, lot GR3190550-20, 1:500 dilution for immunostaining); Rabbit anti-Par-3 (Millipore 07-330; RRID:AB_2101325; lot 3322358, 1:500 for immunostaining); Guinea pig anti-DLIC1-Cter (a gift from Dr. T. Uemura, 1:100 for immunostaining) ^42^; Rabbit anti-Pcm1 (Rabbit antibodies raised against C-terminus of human PCM-1 comprising nucleotides 4993–6095, 1:200 for immunostaining) was provided by Dr. Merdes ^21^.

### Generation of Chicken anti-PCM1 antibody

The human PCM1 C-terminal fragment was cloned into pGEX-2TK GST fusion vector (GE Healthcare Life Sciences) using the primers: 5’-ATA GGA TCC CTG AAA GAC TGT GGA GAA GAT C-3’ and 5’-ATG AAT TCG ATG TCT TCA GAG GCT CAT C-3’ to add the BamHI and EcoRI restriction enzyme sites on both sides of the coding sequence. The plasmids were transformed into *Escherichia coli* XL-1 blue strain by electroporation (Bio-Rad) for PCM1-c-term-GST recombinant expression. The recombinant protein from the cell lysate was purified using Glutathione Sepharose® 4B GST affinity columns (Millpore Sigma, GE17-0756-05). The purified PCM1-GST fusion protein was analyzed on SDS-PAGE stained with Coomassie Blue (Invitrogen). It was used as an antigen for generating IgY anti-PCM1 antibodies produced in the immunized egg yolk by Aves Labs (https://www.aveslabs.com/). The IgY antibodies in the immunized egg yolk were first purified by ammonium sulfate precipitation ^43^, followed by affinity purification on the affinity column (Pierce NHS-Activated Agarose, Thermo Scientific, PO# 26197; Pierce Centrifuge Columns, 10mL, gravity or centrifuge compatible, Thermo Scientific PO# 88988) loaded with PCM1-c-term-GST fusion protein. The purified Chicken IgY anti-PCM1 antibodies were diluted to 1 mg/ml aliquots containing 0.01% NaN3 and stored at -80 °C. The working dilution is 1:100 for fluorescent immunostaining and 1:500 for western blotting.

### Secondary antibodies and DNA labeling

Alexa®-conjugated goat anti-rabbit (Alexa 568, Invitrogen, catalog number: A11011, RRID. AB_143157, lot 792518, 1:2,000 dilution); Goat anti-chicken (Alexa 488, Invitrogen, catalog number: A11039, RRID:AB_142924, lot 2020124, 1:2,000 dilution); Goat anti-mouse (Alexa 488, Invitrogen, catalog number: A11002, RRID:AB_2534070, lot 1786359, 1:2,000 dilution); Goat anti-guinea pig (Alexa 488, Invitrogen, catalog number: A11073, RRID:AB_2534117, lot 46214A, 1:2,000 dilution) or Donkey anti-guinea pig (Alexa 647, Jackson Labs, catalog number: 706-605-148, RRID:AB_2340476, lot 102649-478, 1:2,000 dilution); Anti-Mouse-IgG-Atto647N (Sigma-Aldrich, catalog number: 50185, 1 mg/mL); DAPI solution (1 mg/mL) (Thermo ScientificTM, catalog number: 62248, 1:2,000 dilution).

### Immunocytochemistry

Samples were first washed and pre-incubated in PBS containing 0.1% Tween 20 or 0.25% Triton X-100 (PBS-T; pH 7.4) with 1% DMSO and 5% natural goat serum at 4°C overnight or longer. They were then incubated with primary antibodies in the preincubation solution (PBS-T with 5% natural goat serum) overnight at 4°C. According to the primary antibodies applied, the samples were then washed thoroughly with PBS-T five times × 10 min each time, followed by incubation in Alexa-conjugated goat anti-rabbit (Alexa 568), goat anti-chicken (Alexa 488), goat anti-mouse (Alexa 647), or goat anti-guinea pig (Alexa 647) secondary antibodies (diluted 1:1000) in the preincubation solution for over 2 hours at room temperature or overnight at 4°C. The samples were washed with PBS-T twice for 10 min each, thrice with PBS for 10 min each, and once with 50% glycerol in PBS for 1 hour, followed by infiltration overnight in 80% glycerol/PBS before imaging. Imaging was done using a confocal microscope (Nikon CSU-W1 Spinning Disk/confocal microscopy) with a 100 × oil immersion objective. The z-step of the imaging stack is 0.26 μm.

### Label Retention Expansion Microscopy (LR-ExM)

LR-ExM was carried out as previously described ^31^. In brief, after primary antibody incubation as described above, the cryosections of 1 dpf zebrafish embryos (10 μm thickness) were incubated with trifunctional linkers, NHS-MA-Biotin conjugated anti-Chicken IgG (2 mg/ml) and NHS-MA-DIG conjugated anti-Rabbit or anti-Guinea Pig IgG (2 mg/ml), in the blocking buffer overnight at 4°C in the dark. After washing in 1 x PBS 4 times, 5 min each in the dark, add 40 μL of freshly prepared gelation solution (190 μL of Monomer solution with 5 μL of 8% TEMED and 5 μL 8% APS) to cover the whole tissue sample on the glass. The gelation solution was freshly prepared by deoxygenizing the gel monomer solution (Sodium acrylate 0.86 g/mL, Acrylamide 0.25 g/mL, N,N′-Methylenebis-acrylamide 0.015 g/mL, Sodium chloride 1.17 g/mL in 1× PBS) using a vacuum pump for over 15 min, before adding TEMED and APS, to enhance the effects of trifunctional linkers. Protect the sections from light, incubate in a humidity chamber, and allow to undergo gelation at 37°C for 1 h. Incubate the gel embedded samples in the digestion buffer (8 units/mL of Proteinase K in 50 mM Tris pH 8.0, 1 mM EDTA, 1M NaCl, 0.5% of Triton X-100) on the slides for 4 h at 37°C or overnight at room temperature. At least 10-fold excess volume of digestion buffer was used (> 5 mL for each slide). Tissue sections embedded in gels would slide off the glass surface after sufficient digestion incubation. Wash the gel embedded samples with excess volume of 150 mM NaCl in 6-well plates, with at least 5 mL in each well for 4 times, 20-30 min each time. After washing off the digestion buffer, incubate the gel samples in the post-digestion staining buffer (10 mM HEPES, 150 mM NaCl in MilliQ water, pH 7.5) with 3-5 μM Alexa Fluor® 488-Streptavidin, 3-5 μM goat anti-Digoxigenin/Digoxin Dylight® 594, anti-mouse Atto647N (1:500), and DAPI (1:1,000) for 24 hours at 4°C in the dark. Wash the gel embedded samples 4 to 5 times with Milli-Q water (30 min for each wash) at 4°C in the dark. The samples expand to approximately four times their original size in the last washing and are then ready for imaging with Nikon CSU-W1 Spinning Disk/confocal microscopy by using 60 × water immersion objective. The z-step of the imaging stack is 0.26 μm.

### Secondary antibodies used for tri-functional linker conjugation of LR-ExM

Goat anti-Guinea Pig IgG (H+L) unconjugated secondary antibody (Invitrogen, catalog number: A18771, RRID:AB_2535548); Goat anti-Rabbit IgG (H+L) Cross-Adsorbed unconjugated secondary antibody (Invitrogen, catalog number: 31212, RRID:AB_228335); Goat anti-Chicken IgY (H+L) unconjugated secondary antibody (Invitrogen, catalog number: A16056, RRID:AB_2534729); NHS-MA-Biotin conjugated anti-Chicken IgY, NHS-MA-Biotin conjugated anti-Rabbit IgG, and NHS-MA-DIG conjugated anti-Guinea Pig IgG (Gift from Dr. Shi Lab) ^12^.

### Western blotting

∼ 20 embryos of 28 hpf were dechorionated. After removing the yolk with syringe needle, embryos were washed once with PBS buffer, followed by homogenization in 80 μl SDS sample buffer, the recipe of which can be found in THE ZEBRAFISH BOOK ^44^. Pipetting over 50 times was applied till the lysate is not stringy anymore. The sample tube was transferred into water bath and boiled at 99 °C for 5 ∼10 mins, followed by centrifugation for 10 min at 10,000 rcf. The supernatant was transferred to a new Eppendorf tube which can be stored at −80 °C or used for western blotting immediately. 15 μl of homogenized lysate was used for SDS–polyacrylamide gel electrophoresis (Mini-PROTEAN® TGX™ Precast Gels, 10 wells, Bio-Rad). Proteins were transferred to a Hybond nitrocellulose membrane (Thermo Fisher Scientific, Catalog number: 88025) by a semi-dry blotting technique with Trans-Blot® Turbo™ Transfer System (Bio-Rad) and the membrane was incubated with appropriate primary antibody diluted in 5% dried milk in PBST (1 x PBS, 0.05% Tween-20) overnight at 4°C followed by incubation with corresponding HRP-conjugated secondary antibodies in 2% dried milk in PBST. After the HRP-conjugated secondary antibody incubation, the membrane was rinsed in 1 x PBST once and five times in 1 x PBS, 5-min each. The membrane was then developed using the SuperSignal West Dura Extended Duration Substrate (Thermo Fisher Scientific, Catalog number: 34075) and visualized with the LI-COR Western blotting detection system (LI-COR Biosciences).

### Cryo-sectioning

24-hpf embryos were fixed overnight at 4°C in phosphate-buffered saline (PBS) buffer with 4% paraformaldehyde (PFA). Fixed embryos were washed by 1 x PBS for two times, 5 min each time, and incubated in PBS buffer containing 30% sucrose in the falcon tube overnight at 4°C. After embryos sank to the bottom of the falcon tube, they were then transferred to plastic molds. The sucrose buffer was removed before adding OCT (Tissue-Tek) into the mold to cover the embryos. After orienting the embryos to proper positions in the mold, the block was frozen on dry ice. Blocks can be stored at −80°C up to several months. Frozen blocks were then cut into 10 μm or 20 μm sections on a Cryostat (Leica) and mounted on Superfrost Plus slides (Thermo Fisher Scientific). The slides were dried at room temperature for 2 ∼ 3 hours and then stored at −80°C until use.

### 5-ethynyl-2’-deoxyuridine (EdU) labelling

Click-iT™ EdU Cell Proliferation Kit for Imaging Alexa Fluor™ 555 (Invitrogen™; Catalog number: C10338) was used for whole mount labeling of cell proliferation in 20-22 hpf zebrafish embryos. EdU concentration was diluted 1:1000 (20 μM) in the working solution. For EdU pulse-chase experiment, 20 hpf embryos were pulsed in 1 x EdU working solution in the 6-well plate (3 mL/ well) for 2 hours at 37 °C. Following the pulse, embryos were immediately rinsed in the new well with Ringer’s solution three times (5 min each). The EdU pulse-labeled embryos were moved to a new well with egg water and incubated at 28℃ till next morning. After the EdU pulse-chase, the embryos were fixed with 4% PFA and processed for EdU detection according to the manufacturer’s protocol. Other antibody staining was applied thereafter while the samples were protected from light during immuno-staining. For EdU pulse labeling, 22 hpf embryos were incubated with EdU contained working solution 2 hours at 28 °C followed with fixation and EdU staining directly.

### In vivo co-immunoprecipitation (co-IP)

∼ 200 embryos of 1dpf were placed in 1mL modified Ringer’s solution (116 mM NaCl, 3 mM KCl, 4 mM CaCl2, 5 mM HEPES; pH 7.5) containing Protease Inhibitor (Roche, Cat. No. 04693132001) and Phosstop (Roche, Cat. No. 04906845001). After incubation at room temperature on the shaker for 10 minutes, supernatant was removed and 0.3 ml lysis buffer (50 mM Tris pH 7.5, 150 mM NaCl, 1 mM EDTA, 10% glycerol, 1% Triton X-100) with Protease Inhibitor and Phosstop was added. The tube was kept on ice, and the embryos were homogenized manually by pumping through the syringe with 22-gauge needles for 30-40 times. After homogenization, the tube was placed on the shaker in ice for over 30 minutes. Then the tube was centrifuged for 30 minutes at 10,000 g at 4 °C, and the supernatant was transferred to fresh tube for the following experiment. SureBeads^TM^ Protein A Magnetic Beads were used for co-IP with anti-Pcm1 antibody. The beads were thoroughly resuspended before use; 150 μl beads were transferred to 1.5 ml tubes. After washing with 1ml PBS-T (PBS + 0.1% Tween 20), 200 μl PBS-T with 5 μg antibody was added to resuspend the beads. After rotating for 2 hours at room temperature (RT), magnetized beads were collected, and supernatant was discarded. The antibody-conjugated beads were then washed with 1ml PBS-T for three times and added to the 300 μl embryonic lysate prepared above. After incubation on the rotor at 4 °C for overnight, magnetized beads were collected and supernatant was discarded, followed by five times wash with 1ml PBS-T. Before the last magnetization, the resuspended beads were transferred to a new tube and centrifuged for 30 seconds. Beads were then collected on a magnet and residual buffer was aspirated from the tube. 60 μl 1x Laemmli buffer (diluted by mixing same volume of 1 x PBS with 2x Laemmli buffer, Bio-Rad, Catalog number: #1610737EDU) was added to the beads and incubated for 10 min at 70°C. Eluents were then transferred to a new tube. Final collected samples were boiled at 99°C for 10 min before running SDS-PAGE, or storage at −20 °C.

### Bulk RNA-Seq and analyses

Wild-type AB strain zebrafish (*Danio rerio*) were maintained at 28.5 °C on a 14 hour light/10 hour dark cycle. At the time of mating, breeding males and females were separated in the crossing tank overnight before letting them spawn naturally in the morning for synchronizing developmental stages. Fertilized eggs were grown at 28.5 °C for 24 hours. Each sample was pooled with 12 embryos (in total ∼ 2-3mg) from the same crossing pair. The pcm1 KO mutant with typical morphological defects as pcm1 MO morphants were selected. Three different samples from three independent experiments were used for both WT and KO group of samples. Each pooled sample of embryos were washed by 1ml of PBS at room temperature twice quickly. After removing the liquid from the tube, the sample tubes were frozen in liquid nitrogen immediately. The frozen samples were sent to Genewiz (Genewiz from Azenta Life Sciences) to continue with the standard RNA-Seq, Illumina HiSeq, 2 x150 bp configuration, single index, ∼350M raw paired-end reads per lane, Quality Guarantee is ≥ 80% of bases ≥ Q30. Raw data have been in FASTQ format.

For RNA-seq data analysis, the quality control was first carried out on the raw reads using FASTQ with default parameters, including sequence quality, GC content, adaptor content, overrepresented k-mers, and duplicated reads analysis, aiming at detecting sequencing errors, contaminations, and PCR artifacts. The qualified reads were aligned to the reference genome GRCz11.101 by Hisat2, and the transcripts and corresponding read counts were obtained subsequently. The aligned bam file quality control was achieved using the module bam_stat.py. The count matrix of the transcripts was generated by HTSeq, and the differentially expressed genes (DEGs) were identified by using the module DESeq2. Following the general protocols, a DEG was defined as a gene with log2(FC) > ± 0.5 and p-value < 0.05 between the two groups, *i.e.*, KO vs. WT in this study. The visualization and the analysis of the DEGs were performed via the Pheatmap R package. The molecular functions and the biological pathways of the enriched DEGs were analyzed using the clusterProfiler package ^45^. GO term enrichment analysis was performed with the GO cell component sources, the GO molecular functions sources and the GO biological processes sources in the Metascape ^46^. Enriched terms (P < 0.05, a minimum count of 8, and an enrichment factor of >1.5) were grouped into clusters based on their DE gene similarities and visualized in a network plot using Cytoscape, in which node colors indicate the P value, node sizes indicate DE gene set size, and the connecting edges indicates their DE gene similarities. Processing of RNA-seq data and clustering analysis were performed blindly and unblinded in the following DEG analysis. Heatmaps with hierarchical clustering were made using the heatmap.2 package with the transformed values in which the expression of each gene was scaled across all three samples in both *pcm1* KO and WT groups (z-scored).

### DNA Plasmids, mRNA synthesis and microinjection

Plasmid DNAs (pCS2-H2B-mRFP) were extracted from bacterial clones in stock ^6^. pCS2-par-3-GFP plasmid was a gift from Dr. J. von Trotha ^47^. pCS2-GFP-centrin plasmid was a gift from Dr. W. A. Harris ^48^. The full-length of cDNA of zebrafish Pericentriolar Material 1 (pcm1, Gene ID: 321709, Ensembl: ENSDARG000000-62198) were synthesized and sequenced by Genecreate (www.genecreate.com). The pcm1-linker-GFP plasmids was made by cloning pcm1 cDNA into pcs2-GFP plasmid (gift from the Jan Lab) by Gibson assembly. PCR primers for adding linker between *pcm1* cDNA and GFP cDNA sequence were used as follows: GFP-pCS2-Linker-R: CGT GGA CCC GCC ACC ACT CCC GCC ACC AGA TTT GTA TAG TTC ATC CAT; pcm1-Linker-F: TCT GGT GGC GGG AGT GGT GGC GGG TCC ACG ATG GCA ACG GGT GGC ACT; pCS2-PCM1-R: CCG TTG CCA TGG TGG ATC CTG CAA AAA GAA C; pcm1-pCS2-F: AGG ATC CAC CAT GGC AAC GGG TGG CAC TCC; pcm1-Linker-R: CGT GGA CCC GCC ACC ACT CCC GCC ACC AGA TGC ACT CTG GGC TCC AAT; pCS2-Linker-F: TCT GGT GGC GGG AGT GGT GGC GGG TCC ACG ATG AGT AAA GGA GAA GAA. For par3-GFP, GFP is at the 3’ end of Par-3 protein. For GFP-centrin, GFP is at the 5’ end of Centrin. For pcm1-linker-GFP, GFP is at the 3’ end of Pcm1.

Plasmids (pCS2-H2B-mRFP; pCS2-par-3-GFP; pCS2-GFP-centrin; pCS2-pcm1-linker-GFP) were linearized by the restriction enzyme NotI digestion. NotI - linearized plasmids were purified (QIAquick Gel Extraction Kit) and the 5’-capped mRNAs were synthesized using SP6 mMessenger mMachine kit (Ambion). For mRNA injection, 4 nl mRNAs at 0.5 - 1 μg/μl were mixed with equal volume of injection buffer containing 0.05% phenyl red and injected into the yolk of a 1-4 cell stage embryos. All injections were done with an injector (WPI PV830 Pneumatic Pico Pump) and a micromanipulator (Narishige, Tokyo, Japan).

### Morpholino oligos and microinjections

Knockdown experiments were carried out using previously characterized translational blocking antisense morpholino oligonucleotides (MOs): *pard3ab/par-3* MO (5’-TCA AAG GCT CCC GTG CTC TGG TGT C-3’) ^12^; *pcm1* MO (5’-TGG AGT GCC ACC CGT TGC CAT GAT G-3’) ^49^; Standard control MO (5’-CCT CTT ACC TCA GTT ACA ATT TAT A-3’) was used as injection controls. All MOs were ordered from Gene Tools (https://www.gene-tools.com/) and stored at 300 mM in distilled water. For microinjection, ∼ 4 nL MO at 100 mM was injected into the yolk of 1-2 cell stage embryos containing 0.05% phenyl red (corresponding to 4 ng/zygote).

### In vivo time-lapse imaging, image processing and analyses

All the confocal imaging stacks were captured by Nikon CSU-W1 Spinning Disk/High Speed Whitefield confocal microscopy by using 40 × water immersion objective (for live imaging) and 100 x oil immersion objective (for immunofluorescent imaging) with Micro-Manager 2.0 gamma (μManager, University of California).

For live imaging, the glass-bottom petri dish with embryos was placed on a temperature-controlled stage set at 28.5°C. For *in vivo* time-lapse imaging of internalized Dld, centrin-GFP or pcm1-GFP in dividing RGPs of *Tg [ef1a:Myr-Tdtomato],* z-stacks with 20 to 40 z-planes were acquired consecutively at a 1-μm z-step for each embryonic forebrain region. The exposure time for each fluorescent channel was set at 100 ms by choosing the sequential channel scanning mode for each z-plane. The interval between each z-stack ranged from 12 to 30 s, depending on the z-stack settings of the samples. Usually, 80 volumes of z-stacks were captured for each time-lapse imaging, and the duration spanned around 20 – 30 min.

For imaging HuC-GFP in paired daughter cells, H2B-mRFP mRNA was injected into single cell of *Tg* [*HuC-GFP*] embryos at 16-32 cell-stage. Z-stacks with 50 to 60 z-planes were acquired consecutively with a 1-μm z-step for each volume. The scanning interval between volumes of z-stacks was 6 min. The exposure time for each channel was set at 100 ms for each z-plane as described above. For each embryo we have taken about 200 volumes of z-stacks lasting ∼20 hours. In figure panels and movie frames, maximum intensity projections of 5 to 10 z-planes with 1-mm z-step size were shown, representing the approximate size of RGP and covering both daughters throughout the time-lapsing frames. As for the sequential scanning mode, the frame of different channels might shift a bit due to spontaneous movement of zebrafish embryos, which was later motion corrected during analysis.

After imaging acquisition, Fiji ^50^ was used for image processing. For the measurement of fluorescent intensity of each antibody staining, maximum intensity projection (MIP) of 8-10 z-planes (0.26 μm z-step) was applied to the image stacks to cover the z-planes in the middle of whole RGP. The DAPI staining of each sample was used for the normalization of staining intensities of the other channels of the same sample. The normalized immunofluorescent intensity of the same anti-body was use for the comparison among different samples.

For cell nuclei counting, Fiji plugin for StarDist was used as described ^51^. For the colocalization analyses of Par-3, Pcm1, Rab11 and Dld, the area (50 x 100 pixel, 0.126 μm/pixel) in between the mitotic nucleus was chosen for applying JACoP analyses with Fiji ^31^. For tracking internalized Dld particles in dividing RGPs and determination of distribution at difference mitotic cell phases, Trackmate has been applied for each group of live-imaging dataset as previously described ^12^. For generating the kymograph of internalized Dld dot and plotting the trajectory throughout the mitosis, the spatial registration of each time frame was done by adopting the center point between two centrosomes as the center of dividing RGPs. The two centrosomes labeled with Cen-GFP were used to define the anterior-posterior axis: the anterior centrosome was given the coordinate 0 and the posterior centrosome 1. Each Dld endosome in the RGP cell was then projected onto this axis to obtain its relative distance (value between 0 and 1) at each time frame. On kymograph images, the grayscale value of each pixel indicates the probability of all tracked Dld endosomes at the corresponding location at each time frame. The relative distances of all tracked Dld endosomes were then used for calculating distance and velocity. The temporal registration among different time-lapsing image dataset was done by adopting the anaphase with the first appearance of cleavage furrow to be T=0 min. The asymmetry index of fluorescent immunostaining and live labeling within two newly formed daughter cells was calculated as previously described ^12^. The total fluorescent intensity of antibody immunostaining in both parts of the same RGPs were measured by FIJI and the fluorescent background was set to 0 before the measurement. The asymmetric expression was determined when the total fluorescent intensity of antibody immunostaining in one part is 50% more than the other part (posterior asymmetric when asymmetric index > 0.2 or anterior asymmetric when asymmetric index < -0.2). If less than 50%, the expression was determined as symmetric (-0.2 < asymmetric index <0.2). The asymmetric index of perfect symmetry is “0”, and “1” or “−1” indicates absolute asymmetry (posterior or anterior, respectively).

For LR-ExM images shown in figure panels, Aydin denoising was applied to process the whole stacks of z-planes with all four channels respectively (https://royerlab.github.io/aydin/<u>, DOI 10.5281/zenodo.6612581</u>). “Butterworth” denoising algorithm was used for removing the immunostaining background and noise caused by unspecific or unwastable trifunctional linker signals without removing specific signals.

### Human induced pluripotent stem cell lines

Human induced pluripotent stem cell lines (hiPSC) WTC11 (RRID: CVCL_Y803) and KOLF2.1J (RRID: CVCL_B5P3) were used. All pluripotent stem cell lines were cultured at 37°C on Matrigel (Corning Catalog No. 354277) in Gibco™ StemFlex™ Medium (Thermofisher, Catalog No. A3349401) and amplified using Accutase (Stem Cell Technologies. Catalog No. 07920) for passaging. iPSCs were thawed in the presence of Y-27632 Rock Inhibitor (Stem Cell Technologies. Catalog No. 72304) and the culture medium was changed every day.

### Human iPSC-derived neural rosettes and forebrain organoids

For each independent vial of neural rosette culture, one million human iPSCs were thawed and resuspended with 1 mL of Dulbecco’s Modified Eagle Medium F12 (DMEM F12) media (Thermofisher, Catalog No. 12634010) then diluted in 2 mL of warm DMEM F12 in a 15 mL Eppendorf conical tube (Eppendorf, Catalog No. 0030122151). Our protocol was adapted from previous reports ^34, 52^. hiPSCs were collected by centrifugation at 200 g for 5 minutes at room temperature. After Removal of the supernatant carefully, the cells were resuspended in 2 mL

Gibco™ StemFlex™ Medium (Thermofisher, Catalog No. A3349401) with 10 µM Rock Inhibitor (Stem Cell Technologies. Catalog No. 72304). The resuspended cells (normally > 500k cells) were transferred to a single well on the Matrigel (Corning Catalog No. 354277) pre-coated 6-well plate (Corning. Catalog No.3516). The plate was then placed in the incubator at 37°C and 5% CO2 with saturating humidity. The culture media was changed daily. After incubation for five days, the cultured cells in the well were washed once with PBS. Then 1 mL Accutase was added to cover the cultured cells for 3-5 minutes at 37°C. 5 mL DMEM F12 was added to collect cells by pipetting up and down slowly for ∼ 20 times and transferred to a new 15 mL Eppendorf conical tube. More DMEM F12 media was added to 10 mL total volume. The cells were pelleted at 200g for 5 min at room temperature. The supernatant was removed and the cells were resuspended in 2 mL of differentiation medium N2B27 [Advanced DMEM F12 added with equal volume of Neurobasal (Life Technologies), supplemented with N2 (Life Technologies), B27 without Vitamin A (Life Technologies), penicillin/streptomycin 1%, β-mercaptoethanol 0.1% (Life Technologies)], plus 10 μM Rock-Inhibitor, 0.2 μM LDN (LDN193189, Selleckchem, Catalog No.501362646), 10 μM SB (SB431542, Selleckchem, Catalog No.101762-616) and 5 ng/mL FGF2 (FGF basic 154aa, human, Peprotech, Catalog No.10771-938).

For forebrain organoid cultures, the resuspended cells from above were transferred into a new ultra-low attachment 6 well plates (Corning, Catalog No.07-200-601) with one million cells per well (Day 0). Then the plate was placed back into the incubator and the differentiation media were changed every other day. The rock inhibitor was omitted in the differentiation media. The embryoid bodies (EB) floated up in the medium at day 5. At Day 16 of differentiation, the differentiation media was supplemented with EGF (10ng/mL) and FGF2 (10ng/mL) to promote proliferation and expansion of early organoids. The organoids were collected at day 25 of differentiation. The forebrain organoids with ventricle-like structures were selected for fixation, cryosection, and immunostaining ^34^.

For neural rosette cultures, the resuspended cells from above were transferred into a new low adherent 6-well plate (Fisher Scientific. Catalog No.7200601) with ∼ 300k cells per well. EBs at day 5 were transferred to a new 6-well plate coated with PDL (Millipore. Catalog. MILL-A-003-E)/Laminin (Gibco. Catalog No.2301715) containing fresh media of N2B27 with 0.2 μM LDN and 10 μM SB. The plate was slowly rotated to spread the EB containing media over the well before placing it back into the incubator. The medium was changed once every other day for another 6-8 days. The forming neural rosettes were detected usually starting at Day 12, and collected under a stereomicroscope with a mini cell scraper (Biotium, Catalog No.NC0325221). The neural rosettes were then seeded to a new PDL/laminin pre-coated well containing N2B27 media with 10 ng/mL FGF2, 10 ng/mL EGF (Fisher Scientific. Catalog No. GF144), and 10 ng/mL BDNF (Peprotech, Catalog No.450-10). The plate was placed back into the incubator with a change of the media every other day. After culturing 6-8 days, the neural rosettes were fixed with cold 4% PFA for 5-10 min and processed for immunostaining.

### Processing of forebrain organoids for immunostaining

Forebrain organoids (> 1mm in diameter) were collected, rinsed with PBS, and fixed with 4% PFA for 5 min at 4°C and rinsed with PBS three times for 5 min. The fixed EBs were incubated in 30% sucrose until saturation and sinking to the bottom, followed by embedding in OCT (Tissue-Tek) - filled plastic molds and storage at -80°C. Frozen blocks were cut into 14 µm sections on a Cryostat (Leica) and mounted on Superfrost Plus slides (Thermo Fisher Scientific). The slides were dried at room temperature for 2 to 3 hours and then stored at −80°C until use.

### Statistics and reproducibility

The number of each independent experiment was provided in the figure legends. For immunocytochemistry experiments, multiple sections from individual brain samples were analyzed. For live imaging, three or more dividing RGPs were analyzed from each embryo, depending on the number of mitotic RGPs that were present in each image stack. For each group of samples, sample size was determined to be adequate based on the magnitude and consistency of measurable differences between groups. No randomization of samples was performed. Embryos used in the analyses were age-matched between control and experimental conditions, and sex cannot be discerned at these embryonic stages. Investigators were not blinded to genetically or MO microinjection -perturbed conditions during experiments. Data are quantitatively analyzed. Statistical analyses were carried out using Prism 9 version 9.0.0: the mean value with standard error of the mean (S.E.M.) was labeled in the graphs. The two-tailed unpaired t-test and the *Chi-square* analyses were applied to determine significant differences between groups. Statistical significance is determined as follows: ns, P > 0.05; * P ≤ 0.05; ** P ≤ 0.01; *** P ≤ 0.001; **** P≤ 0.0001.

## Data and materials availability

All data needed to evaluate the conclusions in the paper are present in the paper and/or the Supplementary Materials. Additional data related to this paper may be requested from the authors.

## Code availability

Custom codes written in ImageJ and MATLAB are available from the corresponding authors upon reasonable request.

## Acknowledgment

We thank M. Munchua and E. Lee for excellent animal care; B. Lu, M. Nachury, J. Reiter, and Guo laboratory members for helpful discussions; S. Schmid for helpful comments on the manuscript; D. Larsen, K. Herrington, and UCSF Nikon imaging center for assistance with imaging and data analysis; Dr. X. Shi for sharing all conjugated secondary antibody for LR-ExM; Dr. J. von Trotha for the pCS2-Par-3-GFP plasmid, Dr. W. A. Harris for the pCS2-GFP-centrin plasmid, Dr. A. Merdes for the anti-hPCM1 antibody, Dr. T. Uemura for the anti-DLIC antibody, Dr. Shao and Dr. Gestwicki for the human PCM1 antigen purification. Funding: This project was supported by NIH R01NS120218 and R21 R21NS122053 (to S.G.), the UCSF Mary Anne Koda-Kimble Seed Award for Innovation 2021 (to X.Z.), and Chan Zuckerberg Biohub San Francisco (X.Z. & L.R.).

## Author contributions

X.Z. and S.G. designed the experiments and interpreted the results. X.Z. performed most experiments, Y.W. performed bulk RNA-Seq annotation contributed to Fig. 6, A.C.S. performed kymograph annotation in Fig. 2 and offered guidance for Aydin denoising that contributed to Fig. 5 and Fig. 6, X.C. has provided all codes for live tracking of internalized Dld and results annotation in Fig. 2, V.M. prepared hiPSC cultures used in the study, Z.D. performed CRISPR and assisted full-length zebrafish pcm1 sequencing and clone, L.R. assisted with critical steps in image analysis, and C.J.W. provided reagents. X.Z. and S.G. wrote the manuscript, with the input from all authors.

