## supplementary information for "PCM1 conveys centrosome asymmetry to polarized endosome dynamics in regulating daughter cell fate"

### Supporting Information (Zhao et al.)

#### Extended Data Figures and Figure Legends

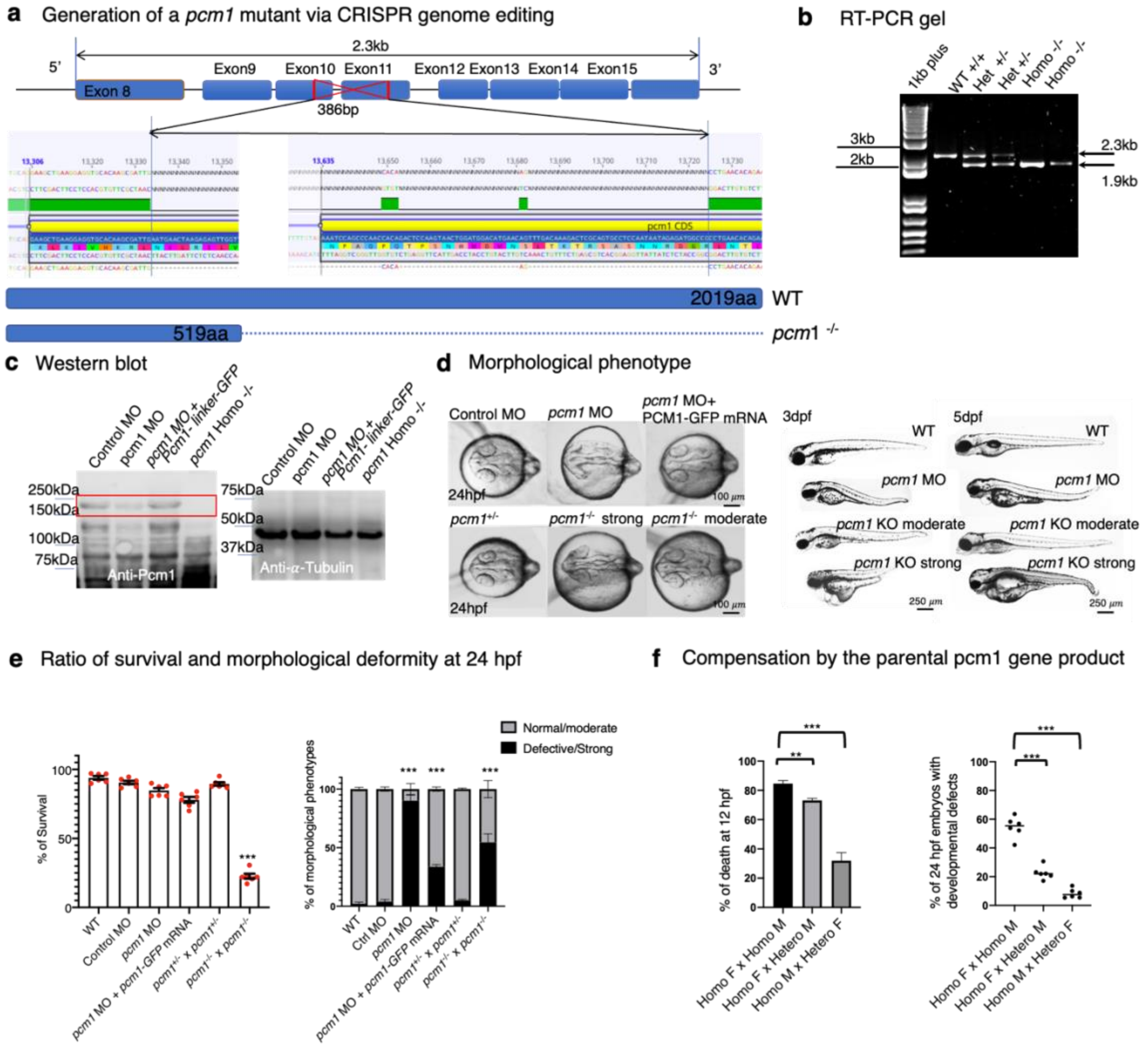


**ED Fig 1. Generation of CRISPR-Cas9 *pcm1* KO and characterization of its developmental defects.**
**a**. A schematic showing the genomic lesion in the *pcm1* locus. The DNA sequence and translated amino acid sequence (Genious Prime 2022) show a 386 bp depletion between Exon 10 and 11 in the *pcm1* KO. The predicted truncated Pcm1 protein is 519 a.a., which is much smaller than WT Pcm1 (2019 a.a.). **b**. Gel electrophoresis of RT-PCR products shows shortened bands from the *pcm1* mutant. The primers flank Exon 8 and Exon 15 as shown in (a). **c**. Western blot with anti-Pcm1 and anti-α-Tubulin antibodies using 1 dpf embryonic extracts. **d**. Dorsal views of 24 hpf embryonic brains (left) and lateral views of 3 and 5 dpf larvae showing the developmental defects of *pcm1* KO mutant compared to control and *pcm1* MO groups. **e**. Statistics of survival and morphological deformity. pcm1 MO and pcm1 KO mutants from homozygous mutant parents showed significantly decreased survival and increased morphological defects at 24 hpf compared to control groups and embryos derived from heterozygous parents. Six independent experiments were performed. *** $p < 0.001$. **f**. Compensation by the parental *pcm1* gene products. Both maternal and paternal *pcm1* gene products contribute to decreased death and developmental defects. Six independent experiments were performed. *** $p < 0.001$; ** $p < 0.01$.

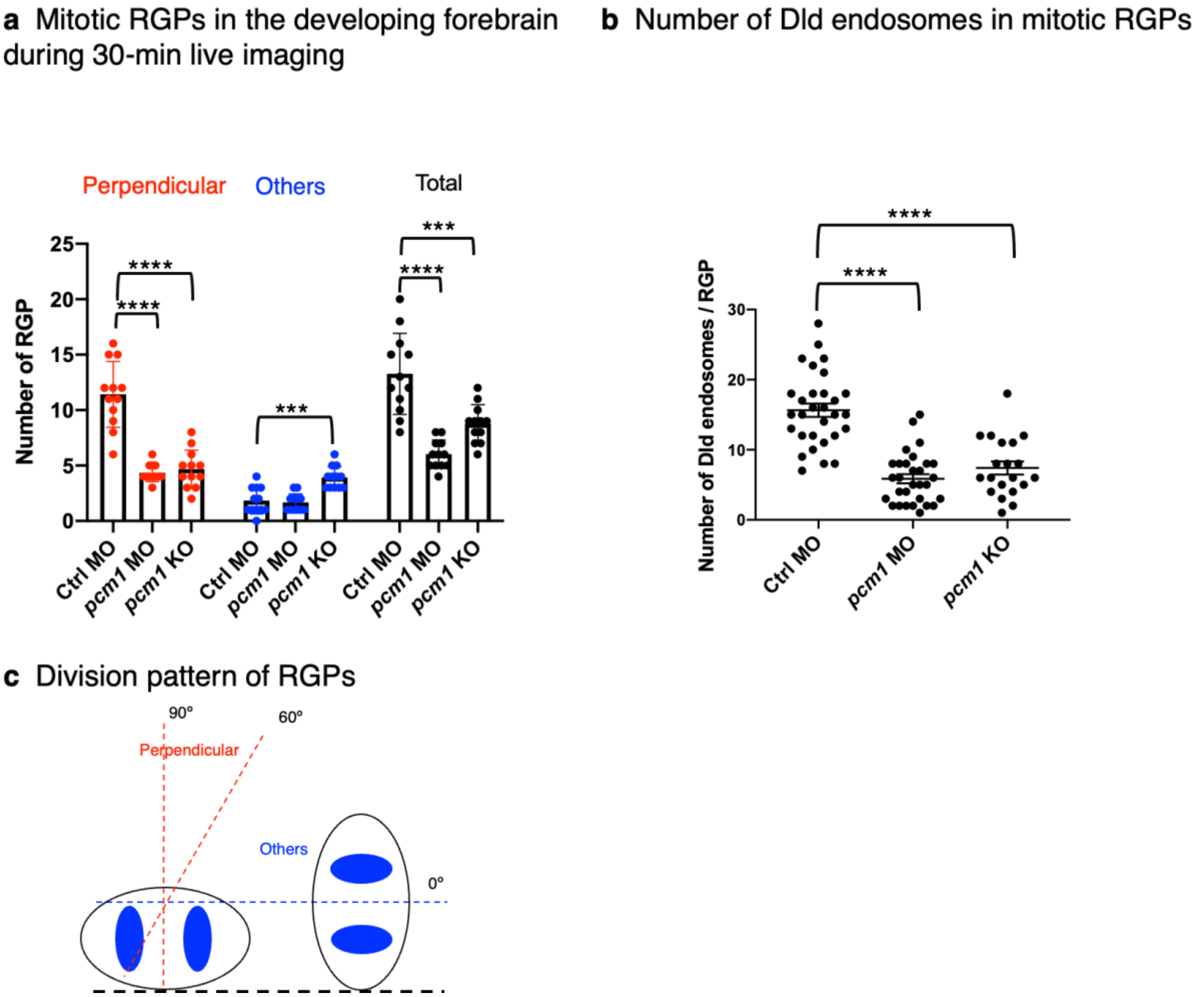


**ED Fig. 2. PCM1 knockdown and knockout decrease mitotic RGPs and Dld endosomes in the developing forebrain.**

**a**. Quantification of mitotic RGPs in the developing forebrain of 24 hpf embryos during 30 min time-lapse imaging. 12 embryos from 4 independent experiments were included in each group. Both *pcm1* MO and *pcm1* KO embryos showed significantly decreased number of mitotic RGPs compared to control. They also showed decreased perpendicularly dividing RGPs and a corresponding increase of non-perpendicularly dividing RGPs compared to control. **** $p < 0.0001$; *** $p < 0.001$; n = 12 embryos per group. **b**. Quantification of internalized Dld endosomes in mitotic RGPs. Mitotic RGPs from both *pcm1* MO and *pcm1* KO embryos showed significantly decreased number of Dld endosomes compared to control. For control MO and *pcm1* MO group, n = 30 RGPs from 6 embryos. For *pcm1* KO, n = 20 RGPs from 6 embryos. **** $p < 0.0001$. **c.** Schematic map of perpendicularly dividing RGPs (dividing plane is 60º-90º to the ventricular surface) and non-perpendicularly dividing RGPs (others). The black dashed line denoted the ventricular surface.

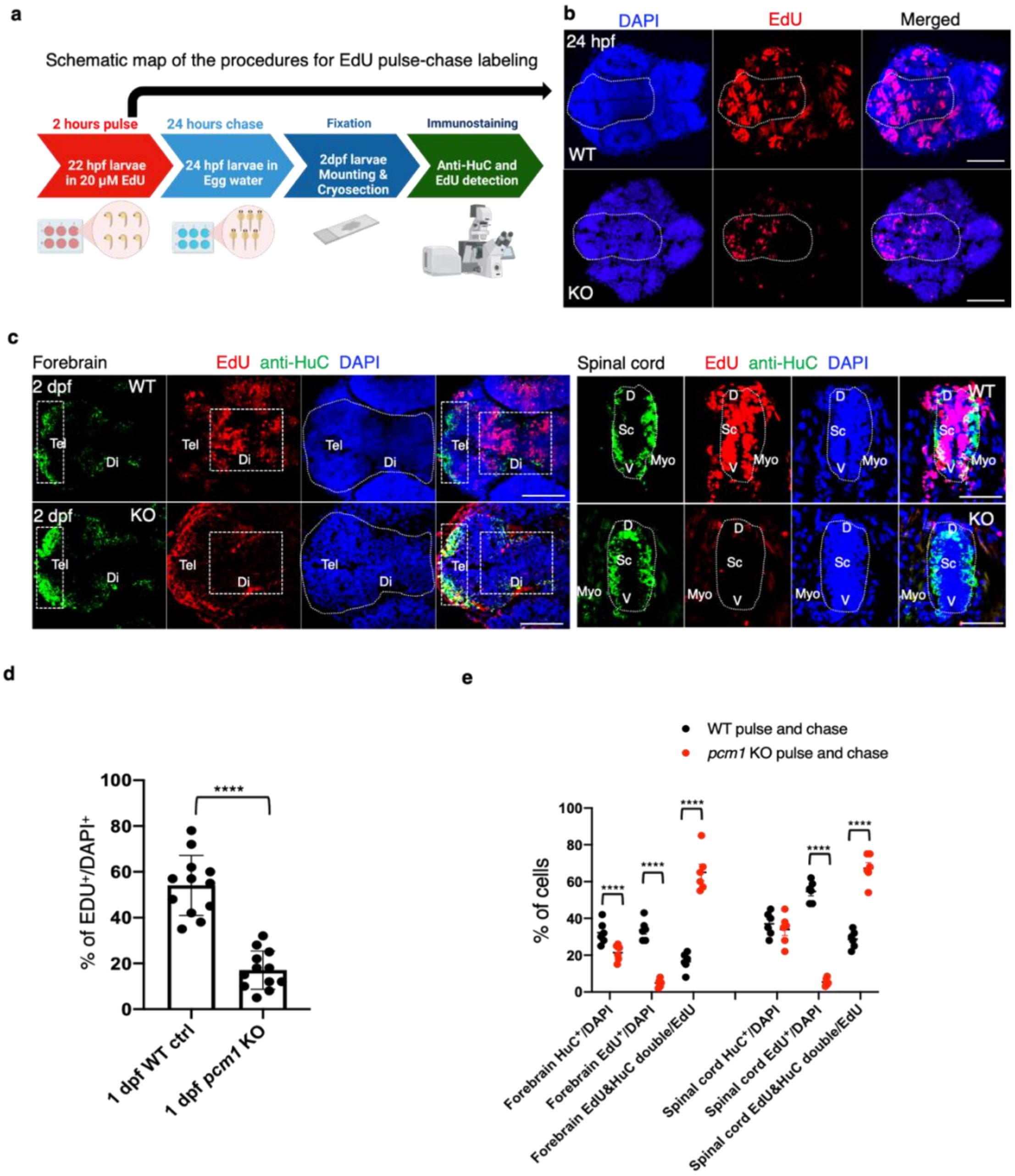


**ED Fig. 3. Pcm1 is required for proliferation and maintenance of progenitors revealed by EdU pulse and pulse-chase labeling. a**. Schematic of EdU pulse only and EdU pulse-chase experiments. **b**. Immunofluorescent images of cryo-sectioned 24 hpf embryonic forebrain following EdU pulse labeling. Scale bar, 100 μm. **c**. Immunofluorescent images of cryo-sectioned 2 dpf embryos (forebrain, left; spinal cord, right) following EdU pulse-chase labeling. D, Dorsal; Myo, Myotome, Sc, Spinal cord; V, Ventral. Scale bar, 50 μm. **d**. Quantification of EdU pulse labeling, shows a decreased ratio of $EdU^+/DAPI^+$ cells in 1dpf *pcm1* KO compared to control. Cryosections from 12 embryos were used for each group. **** $p < 0.0001$, unpaired t test, n=12. **e**. Statistics of EdU pulse-chase labeling from 2 dpf cryosections of forebrain and spinal cord. Cryosections from 6 embryos were used for each group. **** $p < 0.0001$, unpaired t test, n=6.

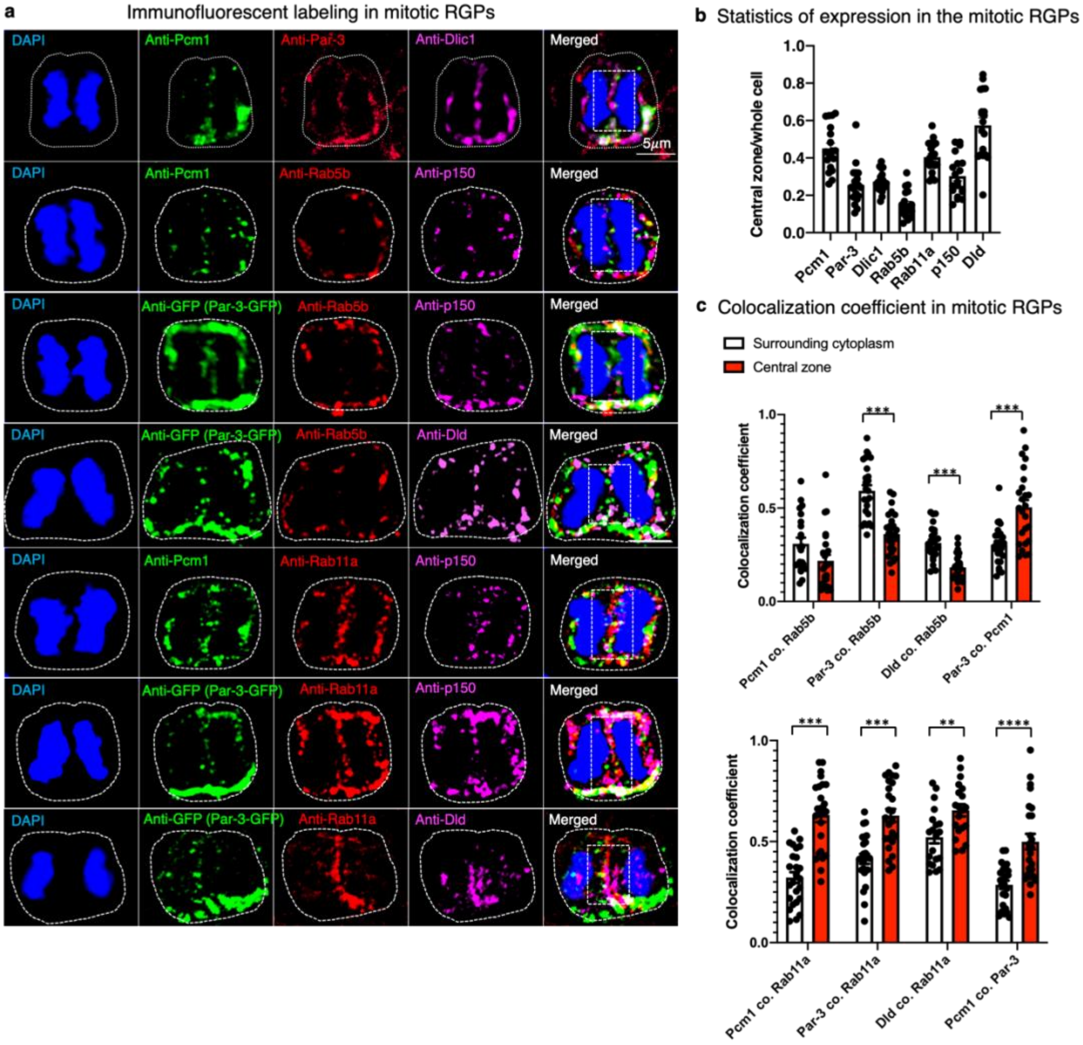


**ED Fig 4. Pcm1 colocalization with Par-3, endosomal, and dynein/dynactin components in anaphase RGPs revealed via conventional fluorescent microscopy and Jacop analyses.** **a**. Immunofluorescent images of Pcm1, Par-3 (or Par-3-GFP), Dld, Rab5b, Rab11a, Dlic1 and p150 in anaphase RGPs from 24 hpf embryonic forebrain. Each image is the maximum intensity projection (MIP) of 10 z-planes. The z-step is 0.26 μm. Scale bar, 5 μm. The area of the whole cell and the surrounding cytoplasm area is outlined by white dashed lines. And the outlined central rectangle (30 x 60 pixels, 1 pixel = 0.126 μm) indicates the central zone in each cell used for the analyses afterwards. **b**. Statistics of protein expression in the central zone of anaphase RGPs. Pcm1, Rab11a and Dld showed over 40% of total cell expression in the central zone, and Par-3, Dlic1 and p150 showed 20% ~ 30% of total cell expression in central zone. Rab5b showed less than 20% of total cell expression in the central zone. 20 cells were included in each group. **c**. Statistics of colocalization coefficient in the different zones of anaphase RGPs. From both charts, Pcm1 showed significantly higher colocalization with Rab11a, but not with Rab5b, in the central zone than in the surrounding cytoplasm. Par-3 and Dld showed significantly higher colocalization coefficients with Rab5b in the surrounding cytoplasm, and higher colocalization coefficients with Rab11a in the central zone. Pcm1 and Par-3 showed significantly higher colocalization in the central zone than in the surrounding cytoplasm. Unpaired t test, n=25 for each group. **** $p < 0.0001$, *** $p < 0.001$,** $p < 0.01$; for "%PCM1 co. Rab5", $p = 0.052445$.

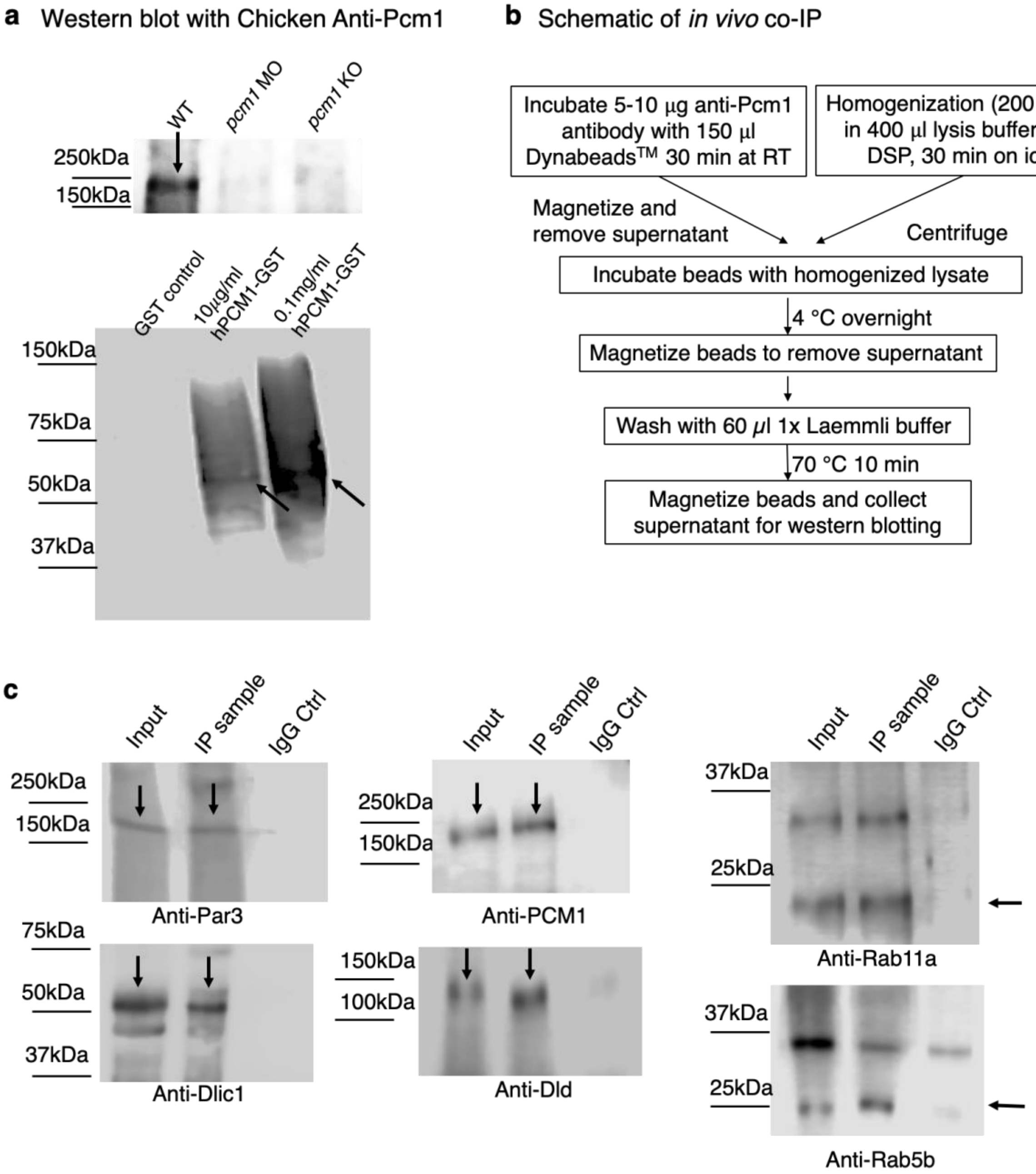


**ED Fig 5. *In vivo* coimmunoprecipitation of Pcm1 with Par-3, endosomal proteins, and dynein components.**
**a**. Western blotting of 1 dpf embryonic lysate (up) and human PCM1 antigen (bottom) with the custom generated chicken anti-PCM1 antibody. **b**. Schematic of *in vivo* Co-IP procedure. **c**. Western blotting of anti-PCM1 co-IPed samples with anti-Par-3, anti-Dld, anti-Dlic1, anti-Rab5b, anti-Rab11a and anti-PCM1 antibodies. Arrows indicate the specific band on each blot.

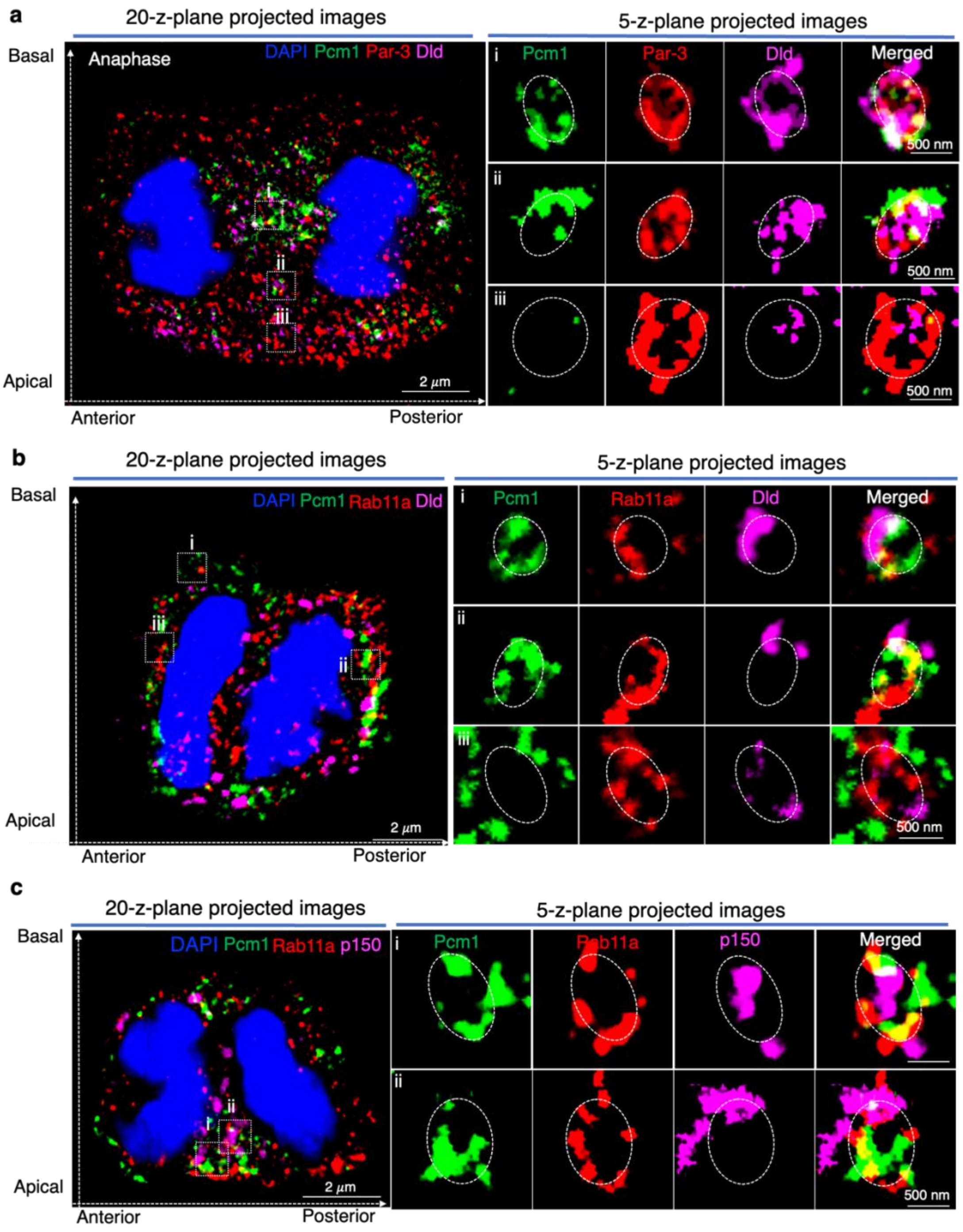


**ED Fig 6. Additiona LR-ExM images of Pcm1 colocalization with Par-3, Rab11a, and P150 on Dld endosomes in developing zebrafish forebrain mitotic RGPs. a**. LR-ExM images of mitotic RGPs immuno-stained with anti-PCM1, anti-Par-3, anti-Dld, and DAPI. **b**. LR-ExM images of mitotic RGPs immuno-stained with anti-PCM1, anti-Rab11a, anti-Dld, and DAPI. **c**. LR-ExM images of mitotic RGPs immuno-stained with anti-PCM1, anti-Rab11a, P150, and DAPI. 20 z plane projected whole cell images were shown on the left, and 5 z plane projected individual Dld endosomes were shown on the right. The z-step is 0.26 µm. Scale bars denote the biological size.

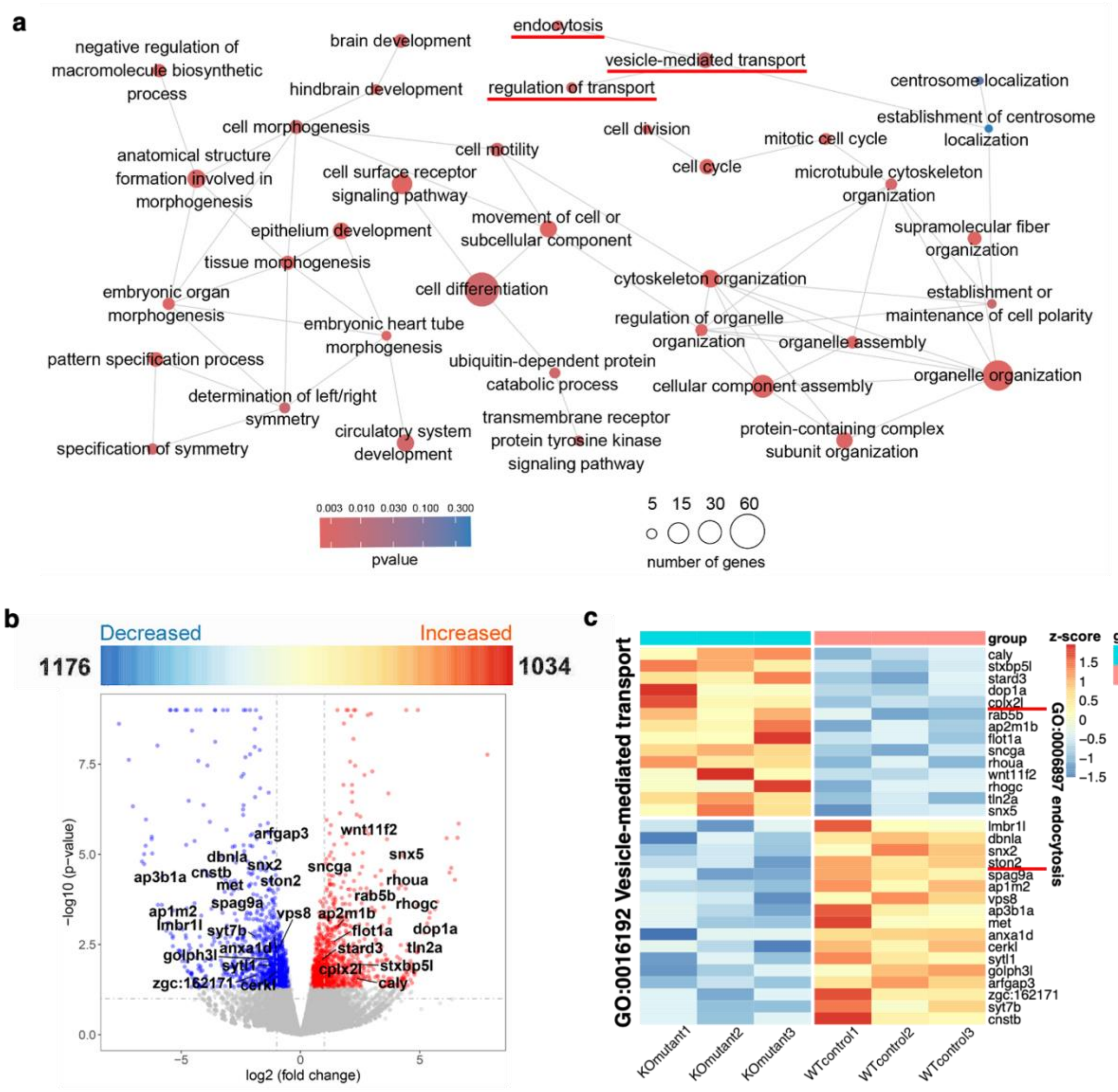


**ED Fig 7. Transcriptomic profiling of *pcm1* KO embryos uncovers dysregulation of genes involved in endocytosis and brain development.**

**a**. A diagram showing a network of 37 significantly enriched GO terms of differentially expressed genes (DEGs) in *pcm1* KO embryos compared to control, built using the Cytoscape Enrichment Map ($p < 0.05$, minimal gene set size = 8). Each node represents a GO term with the color indicating P values. The size of each dot indicates the number of DEGs (5-60) for that specific GO terms. The connecting edges indicate their membership similarities. Two centrosome-related GO terms (colored in blue) were shown but they did not reach the statistical threshold of 0.05, suggesting that the role of Pcm1 in centrosomal regulation is largely non-transcriptional. **b**. Volcano plot of all genes significantly increased or decreased in *pcm1* KO compared to WT ($p < 0.05$, n = 3). The genes involved in vesicle transport, neurogenesis, and cell mitosis are shown on the plot map. **c**. Heatmap showing the relative expression of 31 DEGs involved in endocytosis in *pcm1* KO and WT groups (z-scored).

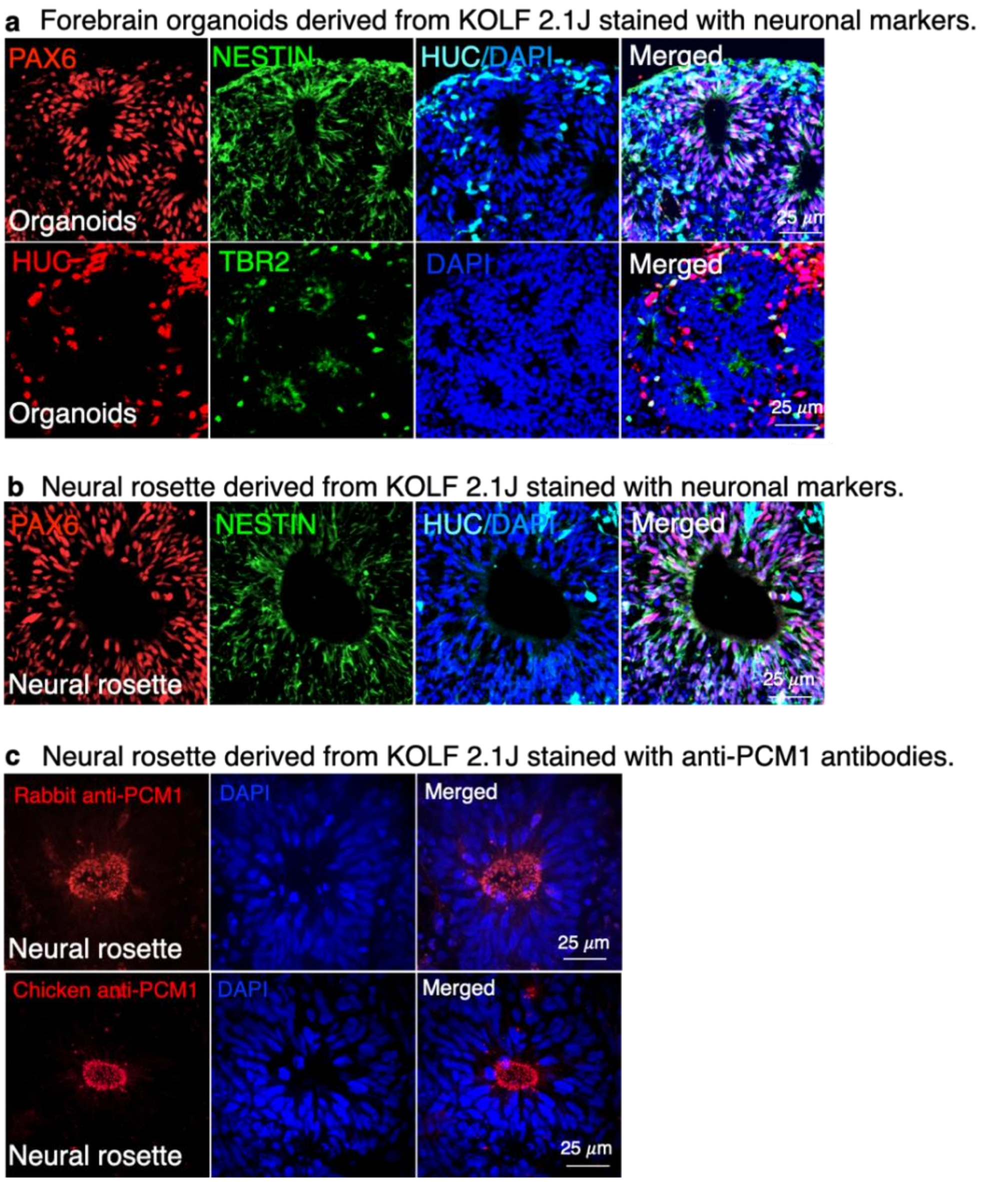


**ED Fig 8. Characterization of marker gene expression in hiPSC-derived forebrain organoids and neural rosettes. a.** Immunofluorescent staining of cryo-sectioned forebrain organoids derived from the KOLF2.1J hiPSC line, with anti-HuC (neuronal marker), anti-Tbr2 (intermediate progenitor marker), anti-Pax6 (neural progenitor marker), and anti-Nestin (neural progenitor marker). **b**. Immunofluorescent staining of neural rosettes derived from the KOLF2.1J hiPSC line with anti-Pax6, anti-HuC, and anti-Nestin. **c.** Immunofluorescent staining showing our custom chicken anti-PCM1 antibody specificity in comparison with a previously reported rabbit anti-PCM1 antibody. Similar enriched fluorescent staining along the apical layer of neural rosettes were observed.

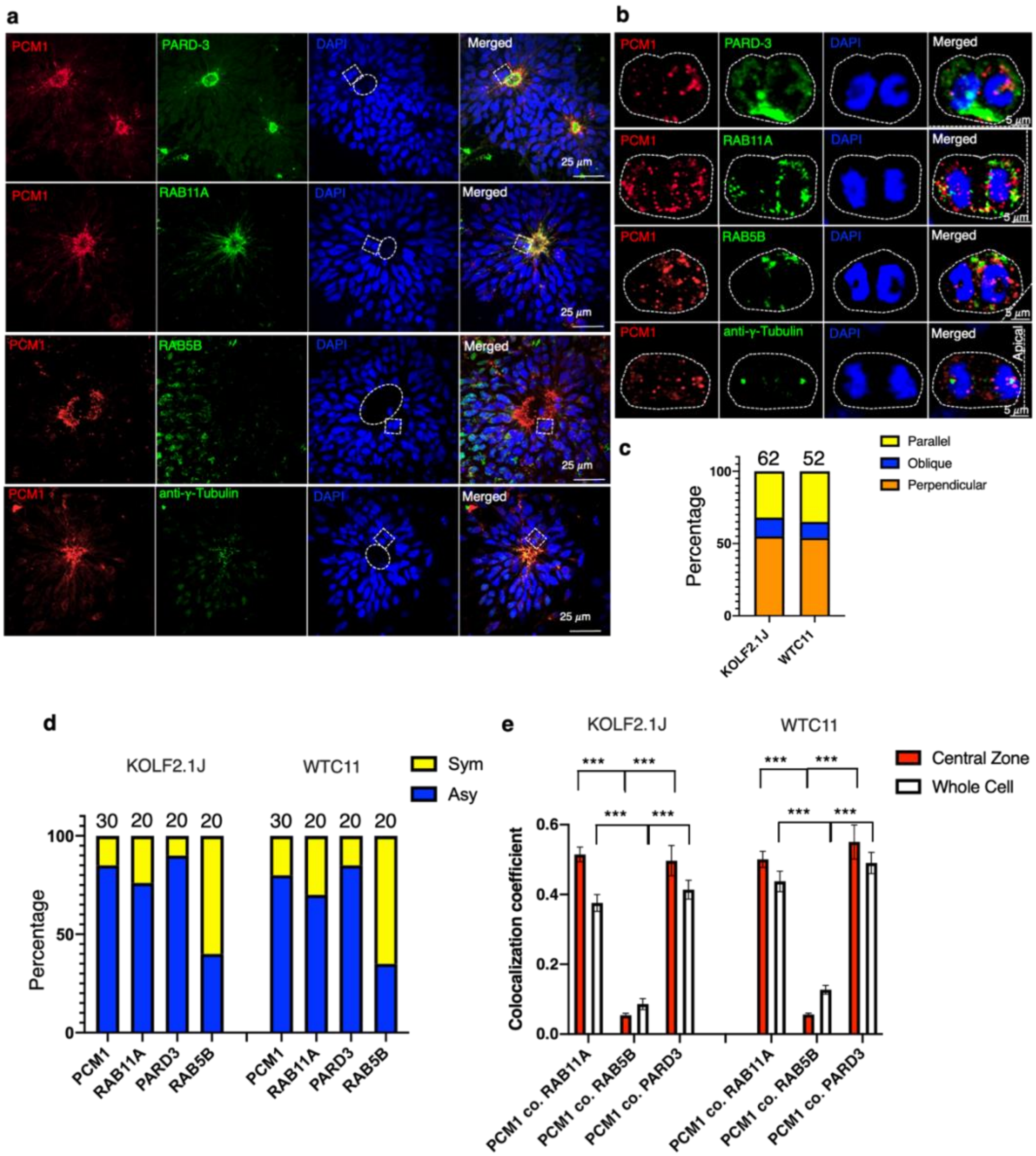


**ED Fig 9. PCM1 expression in mitotic neural progenitors of forebrain neural rosettes derived from hiPSCs.**
**a**. Immunostaining of hiPSC-derived forebrain neural rosettes with anti-PCM1, PARD3, RAB11A RAB5B, and γ-Tubulin. The VZ apical layers were marked with white dashed circles. MIP of 20 z-planes was shown. Z-step is 0.26 μm. Scale bar, 25 μm. **b.** Enlarged views of mitotic NPCs marked by rectangles in (a). MIP of 10 z-planes was shown. Z-step is 0.26 μm. Scale bar, 5 μm. **c.** Statistics of division orientations of RGPs in neural rosettes derived from both KOLF2.1J and WTC11 cell lines. The neural rosettes were from four independent experiments. Ten rosettes from each experiments were selected randomly for statistics. **d**. Statistics of PCM1, RAB11A, PARD3, and RAB5B distribution in NPCs at anaphase. In forebrain neural rosettes derived from both hiPSC lines, PCM1, RAB11A, and PARD3 were asymmetrically distributed in most NPCs. In contrast, RAB5B was symmetrically distributed in most NPCs. **e**. Statistics of colocalization coefficients of PCM1 with PARD3, RAB11A and RAB5B in anaphase NPCs of forebrain neural rosettes. In the central zone, colocalization coefficients of PCM1 with RAB11A (PCM1 co. RAB11A) and PCM1 with PARD3 (PCM1 co. PARD3) are significantly higher than PCM1 with RAB5B (PCM1 co. RAB5B). The colocalization coefficients of PCM1 with PARD3 and RAB11A in the whole cell is less than the colocalization coefficients in the central zone, but still significantly higher than the colocalization coefficients of PCM1 with RAB5b. *** $p < 0.001$, unpaired t test. n=20.

**Supplemental Video Legends**

**Suppl. Video 1-3**. Time lapse recordings of RGPs labeled with Pcm1-GFP (green), Myr-Tdt (red) and Dld (Magenta) in the forebrain of ~24 hpf zebrafish embryo, which were shown in Fig 1h (from top lane to bottom). Each frame is MIP of 8 z-plane (z step is 1 μm) and scanning interval is 30 sec.

**Suppl. Video 4-6**. Time lapse recordings of RGPs labeled with centrin-GFP (green), Myr-Tdt (blue) and Dld (magenta) in the forebrain of ~24 hpf zebrafish embryo, which were shown in Fig 2a. Video 4 is RGP from control MO embryo; Video 5 is RGP from pcm1 MO embryo; Video 6 is RGP from pcm1 KO embryo. Each frame is MIP of 8 z-plane (z step is 1 μm) and scanning interval is 30 sec.

**Suppl. Video 7-9**. Time lapse recording of RGPs labeled with H2B-mRFP (red), Par-3-GFP (green in apical side), in *Tg [HuC-GFP]* zebrafish embryonic forebrain, which were shown in Fig 3d. Video 7 is RGP with P/P division; Video 8 is RGP with P/N division; Video 9 is RGP with N/N division. Each frame is MIP of 10 ~ 15 z-plane (z step is 1 μm) and scanning interval is 6 mins. The time lapse imaging started from 20 hpf and lasted till 36 hpf.
